# A high-quality reference genome for the Ural Owl (*Strix uralensis*) enables investigations of cell cultures as a genomic resource for endangered species

**DOI:** 10.1101/2025.01.22.633903

**Authors:** Ioannis Chrysostomakis, Annika Mozer, Camilla Bruno Di-Nizo, Dominik Fischer, Nafiseh Sargheini, Laura von der Mark, Bruno Huettel, Jonas J. Astrin, Till Töpfer, Astrid Böhne

## Abstract

**Background:** Reference genomes have a wide range of applications. Yet, we are from a complete genomic picture for the tree of life. We here contribute another piece to the puzzle by providing a high-quality reference genome for the Ural Owl (*Strix uralensis*), a species of conservation concern and efforts affected by habitat destruction and climate change.

**Results:** We generated a reference genome assembly for the Ural Owl based on high-fidelity (HiFi) long reads and chromosome conformation capture (Hi-C) data. It figures amongst the best avian genome assemblies currently available (BUSCO completeness of 99.94 %). The primary assembly had a size of 1.38 Gb with a scaffold N50 of 90.1 Mb, while the alternative assembly had a size of 1.3 Gb and a scaffold N50 of 17.0 Mb. We show an exceptionally high repeat content (21.07 %) that is different from those of other bird taxa with repeat extensions. We confirm a *Strix* characteristic chromosomal fusion and support the observation that bird microchromosomes have a higher density of genes, associated with a reduction in gene length due to shorter introns. An analysis of gene content provides evidence of changes in the keratin gene repertoire as well as modifications of metabolism genes of owls. This opens an avenue of research if this is related to flight adaptations. The population size history of the Ural Owl decreased over long periods of time with increases during the Eemian interglacial and stable size during the last glacial period. Ever since it is declining to its currently lowest effective population size. We also investigated cell culture of progressive passages as a tool for genetic resources. Karyotyping of passages confirmed no large variants, while a SNP analysis revealed a low presence of short variants across cell passages.

**Conclusions:** The established reference genome is a valuable resource for ongoing conservation efforts, but also for (avian) comparative genomics research. Further research is needed to determine whether cell culture passages can be safely used in genomic research.

## Background

High-quality reference genomes are rapidly becoming available for many branches of the tree of life (https://www.earthbiogenome.org) [1]. These data are now increasingly used for comparative genomic studies on large evolutionary timescales trying to link phenotypes to genotypes [2]. However, even in genomically and traditionally well-studied groups such as birds, most lineages still lack high-quality reference genome assemblies that would allow for detailed studies of genome evolution.

Typical avian karyotypes are composed of macro-and microchromosomes (but see [3,4]). Compared to macrochromosomes, which are typically between 30 and 250 mega base pairs (Mb) in size, microchromosomes have an average size of 12 Mb, although microchromosomes as small as 3.4 Mb have been observed [5,6]. Despite recent efforts to characterise avian genomes and understand their karyotype evolution, less than 10% of all known bird species have a characterized karyotype [7]. The diploid number of about half of these varies between 78 to 82 chromosomes [1]. Regarding the family Strigiformes (owls), karyotype information is available for 13 % of species [7]. Interestingly, microchromosomes encode half of the genes in birds, although they account for only about a quarter of the genome sequence [5,8]. Moreover, the mutation rate of microchromosomes is significantly higher than that of macrochromosomes [9]. Therefore, avian karyotypes, genome structure and especially the microchromosomes deserve more cytogenetic and molecular attention.

To this aim, we here provide a first high-quality reference genome for the Ural Owl (*Strix uralensis*). This species is one of the largest Eurasian owls, inhabiting the Palaearctic lowlands up to the treeline, mainly in the taiga forest belt over a large uninterrupted range from Scandinavia through Siberia to Sakhalin and the Japanese islands. It also occurs in geographically isolated, mixed and deciduous forests of southeastern and central Europe (southern Germany, Czech Republic, Austria, Slovenia and Poland; partly supported by reintroductions). So far, 11 subspecies have been described from its vast distribution based on differences in size and colouration [10]; although not all of these have been widely accepted [11]. Furthermore, the molecular data at hand (i.e., mitochondrial and nuclear marker genes) do not support morphology-based taxonomic distinctions [12].

Ural Owls are nocturnal hunters of small mammals and birds and usually stay in their territories throughout the year [11,13]. As the Ural Owl is sedentary and nests in hollow stumps or tree holes [14,15], it is affected by ecosystem degradation [16]. Nesting sites have been reduced by intensive logging activities, agricultural use, and forestry management [10]. While globally still considered under the IUCN Red List category “Least Concern”, *S. uralensis* went extinct in Austria, southern Germany, and the Czech Republic in the last century, mainly due to direct persecution [17–19]. Successful reintroductions have taken place in these countries (e.g. [17–19]). These central European reintroductions have restored gene flow between the remaining Alpine and European populations [12,20]. The Ural Owl will likely further be affected by climate change, potentially shifting its range to more northern regions [21] and altering breeding times [22]. Correspondingly, the Ural Owl is a species of conservation measures in the European Union under the <u>EU Birds Directive</u> and <u>Nature Habitats Directive</u>. It is also part of the Bern Convention (<u>Convention on the Conservation of European Wildlife and Nature Habitats</u>).

International trade of all Strigiformes is regulated by the Convention on International Trade in Endangered Species of Wild Fauna and Flora CITES.

Cryobanking, defined as the preservation of viable cells and tissues at ultracold temperatures, typically using liquid nitrogen, is considered paramount in preserving the genetic variability of species, especially those facing population decline as the Ural Owl, to ensure population health and persistence [23,24]. Although some instances have been reported where long-term cell culture generated genetic instability and heteroploidy [25,26], it is still unclear how frequent such a phenomenon is and at which stage of cell cultivation it occurs.

Herein, we generated a reference genome for the Ural Owl as a genomic resource to facilitate further research on this species and on Strigidae more generally. We assess the genome assembly quality and provide a first analysis of its gene content. As a species of potential conservation concern and as a proof of principle, we assessed the application of cell culture to produce sufficient DNA in terms of quantity and quality to allow genomics for species with limited biological material. We investigated mutation as a function of passage number (i.e., the transfer of cells from vessel to vessel). To this end, we obtained a cell culture from the same individual that was genome-sequenced, and cultivated the cell lines until passage 10 and subsequently sequenced replicates of passages 5 and 10 (Figure 1).

**Figure 1:**
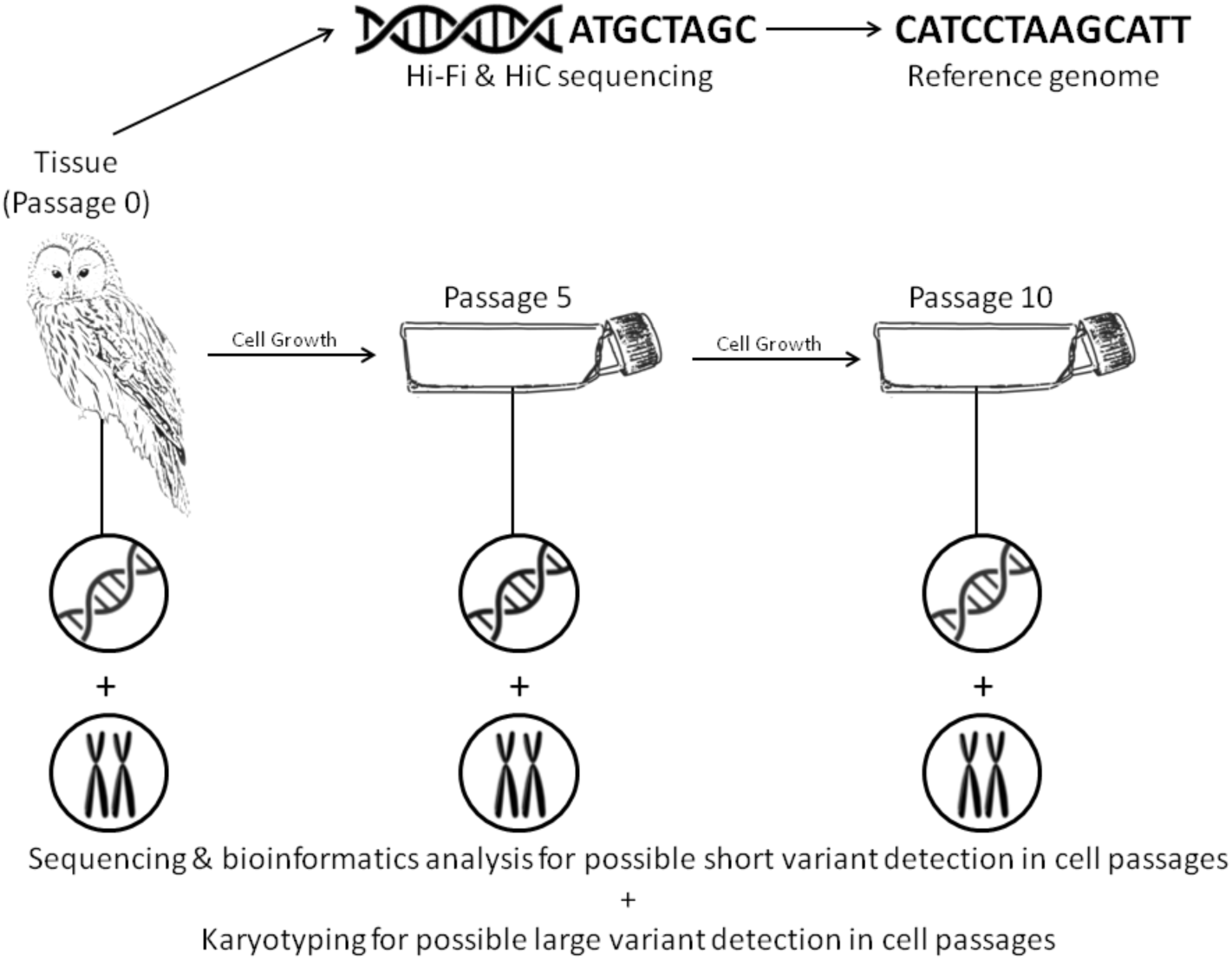
Reference genome and cell passage variant detection workflow. Tissue from a male Ural Owl (*Strix uralensis*) is extracted and sequenced, assembled and annotated to provide a reference genome. Additionally, a cell culture is established from the primary tissue. From passage 0 (primary tissue), passage 5 (three independent replicates) and passage 10 (four independent replicates) cells are harvested for short-read sequencing and karyotyping.

### Data Description

In order to provide valuable genomic resources to the scientific community studying avian ecology and phylogenomics and to investigate the potential of lab-grown cells for use in DNA sequencing, skin cells were harvested from a 10-year-old, recently deceased male Ural Owl individual. The skin samples were, originally, frozen at-80°C and later grown in an appropriate medium and used for DNA sequencing. We performed PACBIO long-read sequencing of muscle tissue, which produces high-quality, long DNA fragments. We used cultured cells for Hi-C sequencing, which allows us to estimate physical proximity of DNA molecules inside the cell to create a bird genome assembly with the highest gene completeness score to date. Next, we grew the harvested skin cells for multiple generations to understand whether this process causes damage to chromosome structure and the accumulation of DNA mutations. In the future, this data can be used to study avian phylogenomics and diversity as well as further understand the unique traits of owls. All sequence data of this study can be accessed from INSDC under the BioProject ID PRJNA1212906. Processed data are available from Zenodo under DOI <u>10.5281/zenodo.14676512</u>.

## Analyses

### Read quality control and estimation of genome size and heterozygosity

After quality control, filtering, and decontamination the final set of HiFi reads used was composed of 5,078,732 reads with a total length of ∼58 Gb and the Hi-C reads used were composed of 79.7 million reads with a total length of ∼ 20 Gb.

Using a k-mer size of 21, GenomeScope was able to predict a genome size of 1,292,799,460 bp, a repeat length of 188,615,362 bp, a heterozygosity of 0.2 % (this would translate to 2 heterozygous sites per 1 kb, a commonly reported heterozygosity indicator for birds) and a read error rate of 0.14 % (Supplementary Table S1). Smudgeplot and GenomeScope both verified the diploid status of the individual. (Supplementary Figure S1; Figure 2). The genome did not reveal any large runs of homozygosity (ROH).

**Figure 2:**
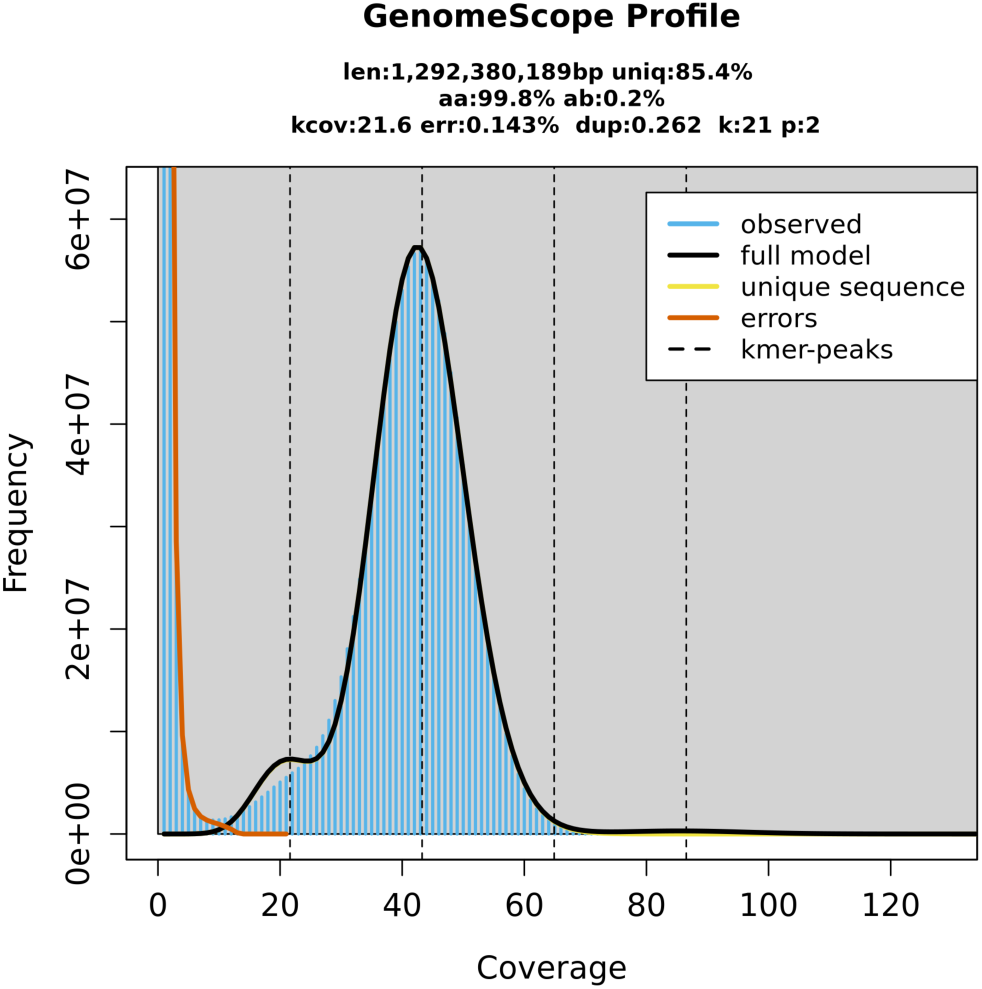
**K-mer genome profile of *Strix uralensis* generated from PacBio HiFi reads with GenomeScope2**.The y-axis shows the k-mer counts and the x-axis shows sequencing depth. The first peak corresponds to heterozygous k-mers and the second larger peak to homozygous k-mers with a coverage of ∼42 x.

### Reference genome

The optimal assembly was created with Hifiasm parameters “-l2--n-weight 5--n-perturb 50000--f-perturb 0.5-D 10-N 150-s 0.4” (Supplementary Table S1).

### Genome Quality Metrics

We could place 93.6 % of assembled scaffolded genome sequence data into 41 chromosomes, which is consistent with the karyotype of the species (Figure 3). We also detected no contamination as all scaffolds aligned to sequences of other avian genomes (Supplementary Figure S2). Our Hi-C contact map further supported the high contiguity of the primary assembly, by showing no remaining conflicts and little to no scaffolds with strong contacts to non-repeat regions (Figure 4, Supplementary Figures S3 and S4).

**Figure 3:**
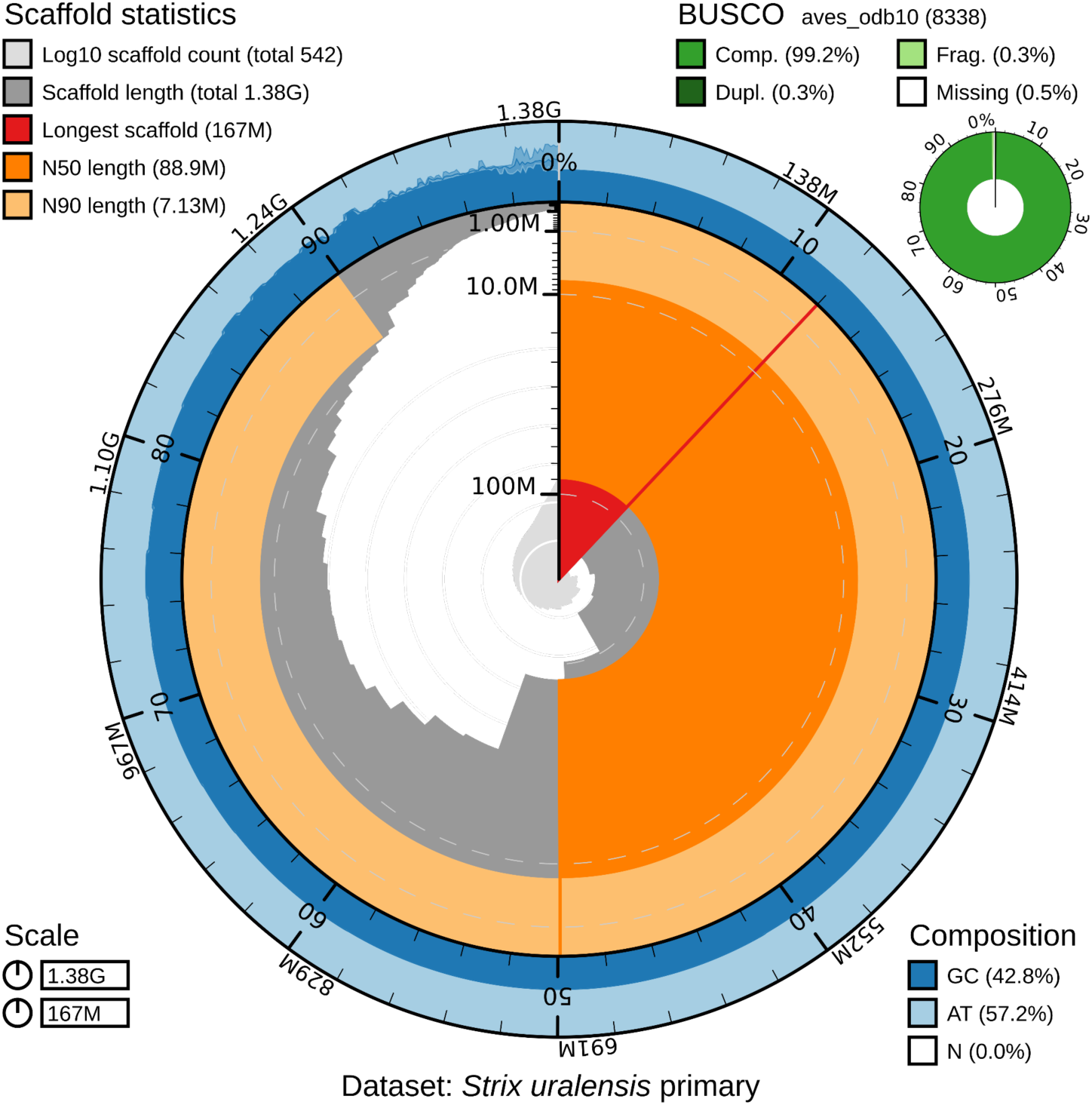
Snail plot summary of assembly statistics for *Strix uralensis* primary assembly. The main plot is divided into 1,000 size-ordered bins around the circumference with each bin representing 0.1% of the 1,381,000,783 bp assembly. The distribution of sequence lengths is shown in dark grey with the plot radius scaled to the longest sequence present in the assembly (166,530,430 bp, shown in red). Orange and pale-orange arcs show the N50 and N90 sequence lengths (88,922,949 and 7,132,230 bp), respectively. The pale grey spiral shows the cumulative sequence count on a log scale with white scale lines showing successive orders of magnitude. The blue and pale-blue area around the outside of the plot shows the distribution of GC, AT and N percentages in the same bins as the inner plot. A summary of complete, fragmented, duplicated and missing BUSCO genes in the aves_odb10 set is shown in the top right.

**Figure 4:**
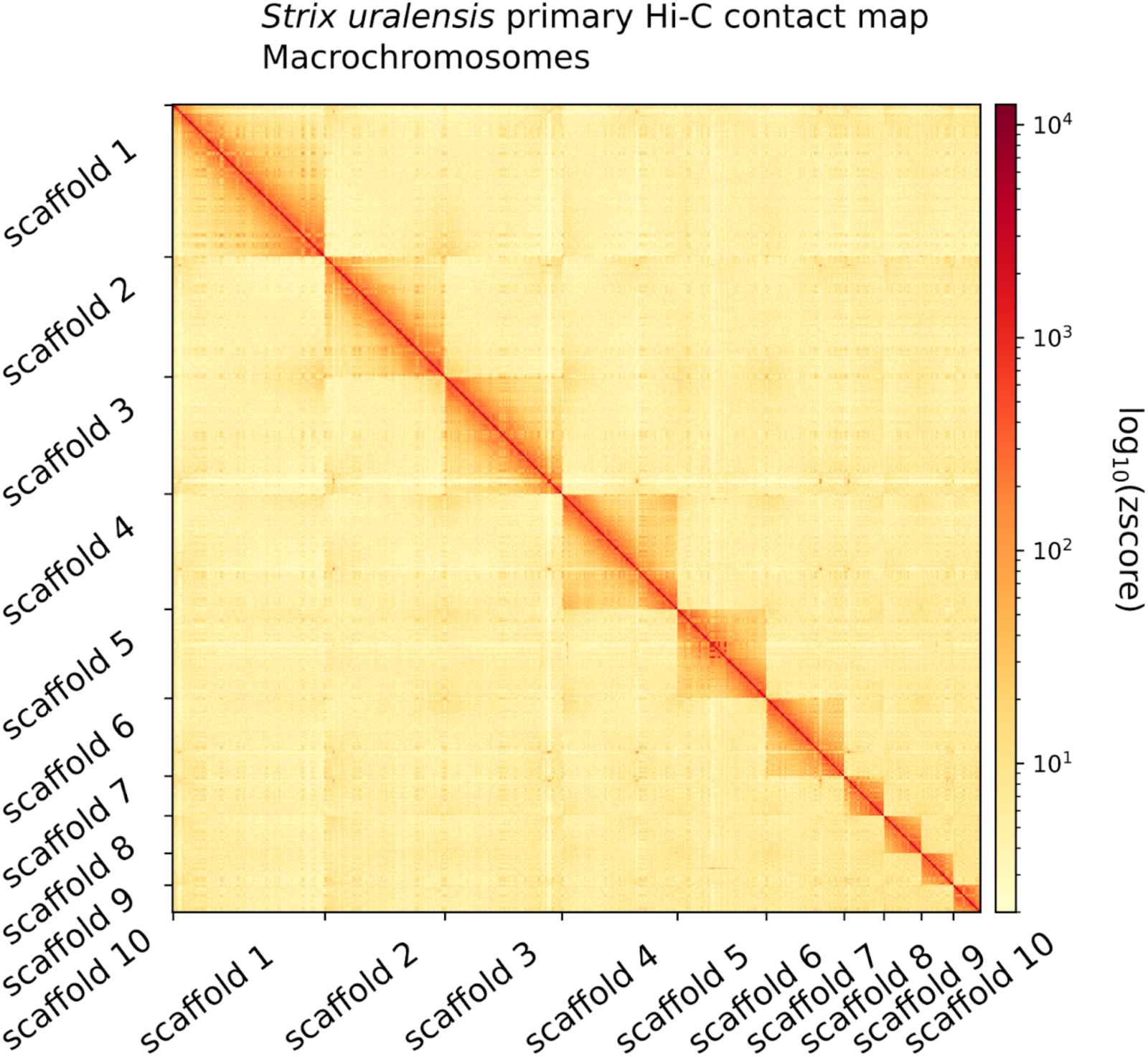
S*t*rix *uralensis* primary haplome Hi-C contact map showing spatial interactions between the ten largest chromosomes. Chromosomes are ordered by size from left to right and from top to bottom. The red diagonal corresponds to intra-chromosomal contacts and depicts chromosome boundaries. The frequency of contacts is shown on a logarithmic heatmap scale. Plot generated with HiCExplorer.

The Merqury Quality Value (QV) score, which is the proportion of the assembly sequence supported by HiFi reads, was estimated for both haplomes. We obtained a score of 64.2 (equivalent to an error probability of 3.8238e-07 %) for the primary and 57.4 (equivalent to an error probability of 1.80919e-06 %) for the alternate assembly (Supplementary Table S2). We also find a completeness score of 98.36 % for the primary and a combined 99.81 % for the two haplomes, representing the fraction of high-quality k-mers from the reads present in the assembly. This further supports the completeness and accuracy of the assembly (Supplementary Table S2).

Aligning the PacBio HiFi, and Illumina Hi-C reads to both haplomes revealed comparable coverage levels (primary: 41.78 ± 12.18, 14.18 ± 91.47-fold respectively; alternate: 34.15 ± 23.24, 12.10 ± 107.37 respectively), mapping rates (primary: 99.85 %, 99.92 % respectively; alternate: 80.52 %, 86.6 % respectively) and mapping quality scores (primary: 36.93, 28.90 respectively; alternate: 28.11, 8.2 respectively). These results further indicate that the assembly is well-phased with a minimal amount of assembly bias and errors (Supplementary Table S3).

**Table 1:** Assembly statistics of the primary and alternate genome assembly of *Strix uralensis*.

| Assembly statistics | Primary | Alternate |
| --- | --- | --- |
| Assembly size [bp] | 1,381,008,983 | 1,262,176,999 |
| GC content [%] | 42.77 | 42.81 |
| Contigs | 512 | 15,615 |
| N50 | 90,173,155 | 17,018,198 |
| L50 | 6 | 18 |
| L90 | 28 | 8,171 |
| Ns per 100 kb | 2.94 | 68.29 |
| Merqury Error (%) | 3.82794e-07 | 1.80919e-06 |
| Merqury QV score | 64.17 | 57.43 |
| Complete BUSCOs [%] | 99.94 | 70.82 |

### Genome Repeat and Gene Annotation

Repeat landscapes depict the clustering of transposable elements (TEs) in relation to their Kimura substitution rates, which measures the divergence of TEs from their respective consensus sequence. Lower Kimura substitution rates indicate recent transposition events, while higher rates suggest older events. From the landscape of the primary haplome (Figure 5, Supplementary Figures S5 and S6), a strong signal for a recent repeat expansion of LINEs (long interspersed nuclear elements) and an even more recent expansion of LTR (long terminal repeat) retrotransposons as well as a well-maintained large number of older repeats are visible. This might support a repeat expansion in the Ural Owl or the genus *Strix*. A third and older expansion is dominated by LINEs and unknown repeats suggesting that they are either a new or unique feature of *Strix* and a reference or consensus might not yet exist in the reference databases.

**Figure 5:**
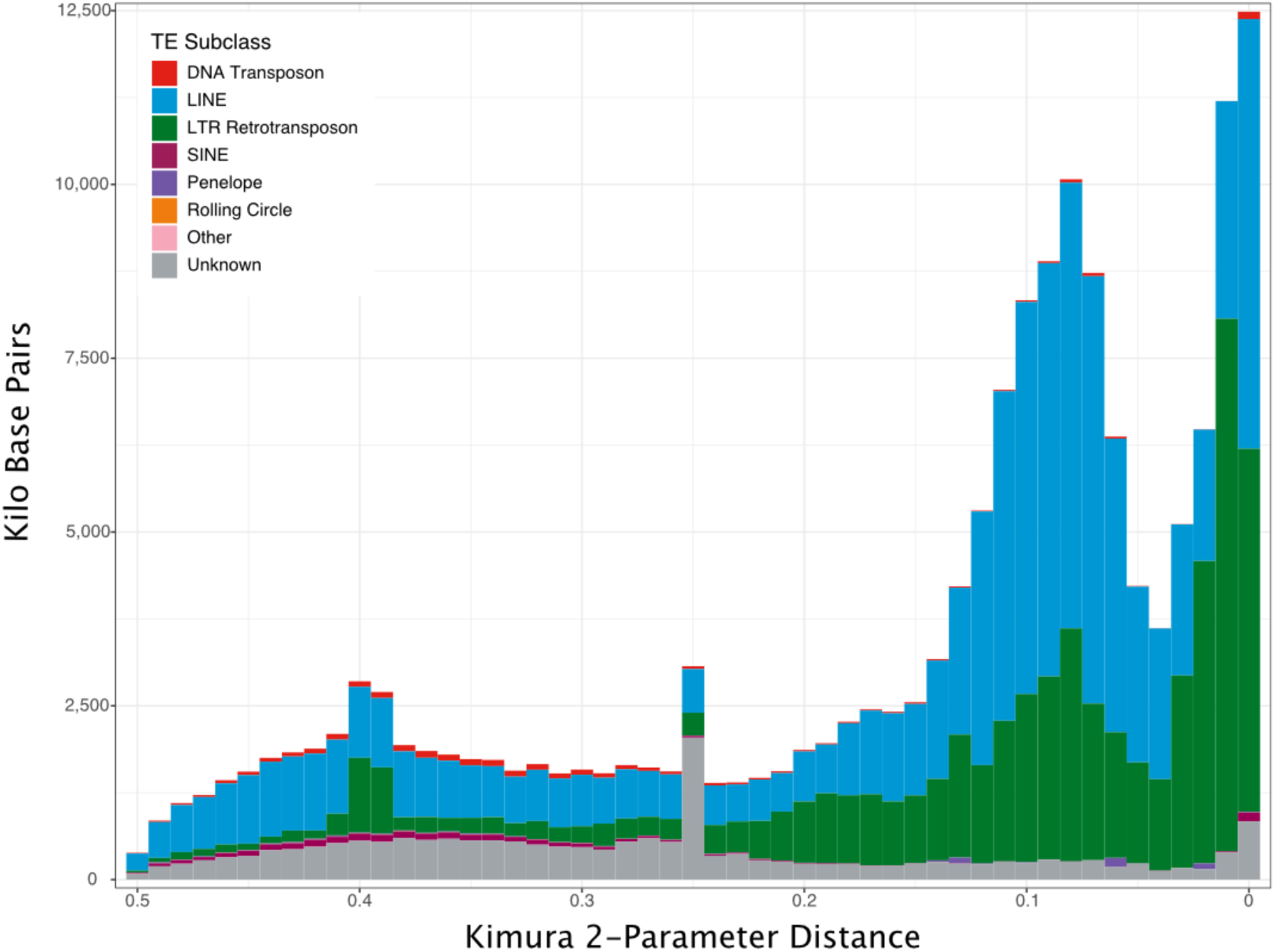
S*trix uralensis* primary haplome repeat landscape. The x axis shows the Kimura substitution of detected repeat categories and the y axis the number of repeats detected for each TE family in kilo base pairs. Detected subclasses are colour-coded as indicated in the inset. The genome assembly was masked using EarlGrey.

### Gene annotation

For the primary haplome, we were able to annotate a total of 17,977 protein-coding genes which cover ∼33.6 % of the total size of the assembly (Supplementary Table S4). We detected 182,313 exons and 164,373 introns. Compared to the Swiss-Prot and UniProt databases we were able to match 16,461 and 17,511 of our genes to annotations respectively (Supplementary Table S5).

We next investigated gene distribution along the genome. Using a 30 Mb cutoff [5,6], we identified ten macrochromosomes and 31 microchromosomes based on our assembly (Figure 4 and Supplementary Figure S7). Despite their size, microchromosomes had a higher gene density than macrochromosomes. While there are comparatively more genes on microchromosomes, these genes are shorter than those on macrochromosomes, mainly due to shorter introns (Supplementary Figure S7).

To shed light on the genome annotation content, we compared the Ural Owl genome to other high-quality genomes of the Aves lineage, including several owl species. Gene expansion (gain) and contraction (loss) among our selected species found 316 gene family gains in the Ural Owl and 207 losses, 168 of which were, presumably, completely lost and, thus, have no representative in the Ural Owl genome assembly (Figure 6).

**Figure 6:**
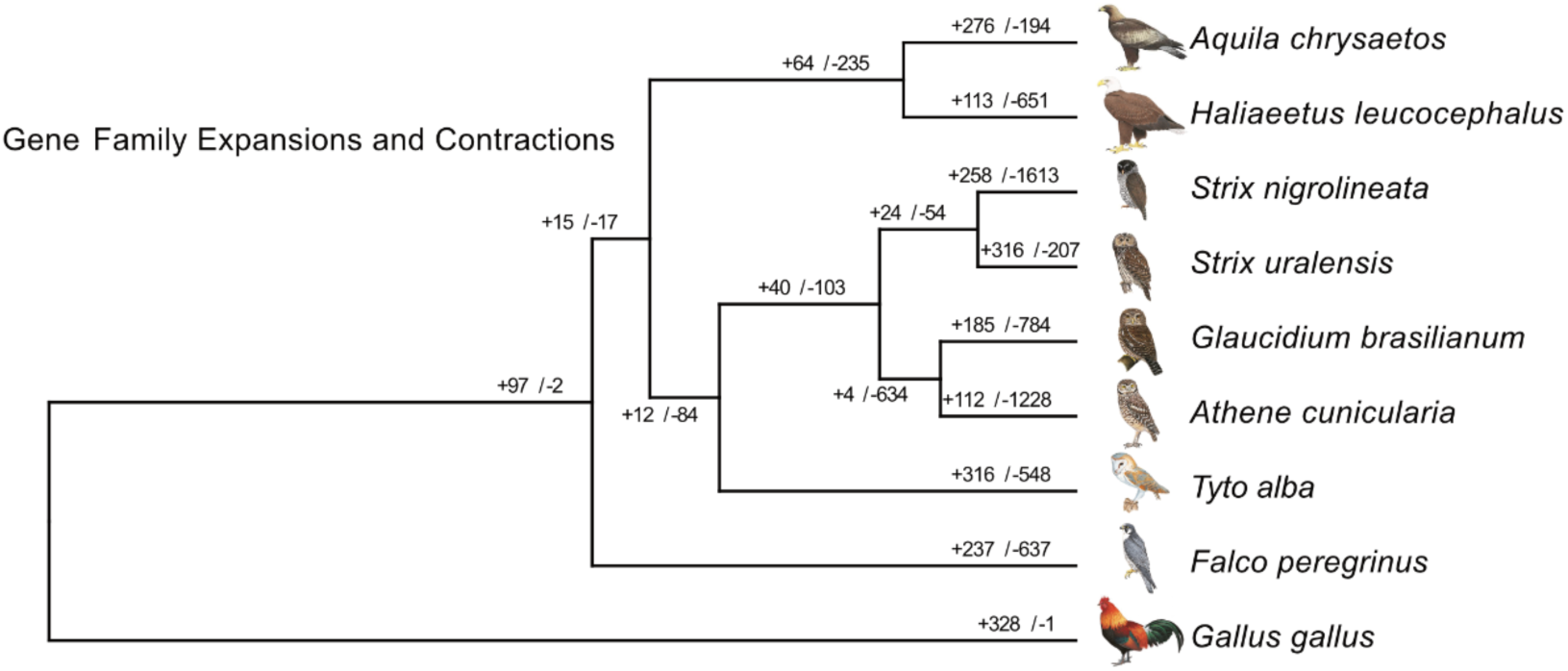
Ultrametric phylogenetic tree of selected Neoaves species and *Strix uralensis*. Numbers indicate gene family expansions (+) and contractions (-). Bird drawings from https://birdsoftheworld.org/.

Additionally, we found 81 gene families unique to the Ural Owl that do not have orthologs in the other species (Supplementary Table S5). We further found that genomes of lower quality, such as those of the Ferruginous pygmy owl, *Glaucidium brasilianum,* and the Black-and-white Owl, *Strix nigrolineata*, had more gene losses, which are hence probably not biologically true but represent technical limitations. A Gene Ontology (GO) term analysis of the genes unique to the Ural Owl revealed many interesting gene families that due to the low quality of the Black-and-white Owl genome might also be interpreted as partially representing the *Strix* genus (Supplementary Figure S8). Among these categories we note several GO terms relevant to characteristic traits of the Ural Owl, namely its adaptation to dim-light conditions and a sedentary and predatory hunting strategy. The “animal organ morphogenesis” parent GO term groups the child GO terms “eye development”, “sensory organ development”, “neurogenesis” and “heart development”, all of which point to adaptations of *Strix*, either to their environment or lifestyle.

Next, we investigated gene gain and loss at nodes that are supported by more than one reference genome which would make them more robust and at the same time informative about clade-specific genomic changes.

We observed 15 gene family gains and 17 losses in the last common ancestor of Strigiformes and Accipitriformes (hawks, eagles, vultures, kites), both characterized by a predatory lifestyle. Overarching GO terms among the gained genes included “behavior”, metabolic, cellular and developmental processes. Notably, the child GO terms contained many terms related to general and cellular metabolism (e.g., “ATP metabolic process”, “carbohydrate derivative metabolic process”, “cellular lipid catabolic process”, “cellular lipid metabolic process”). We identified three gains in keratin genes (feather and scale keratin), two related to histones/histone modification and two related to skeletal muscle functioning (BEST3, CKB).

The gene losses comprised several mitochondrial genes which we attribute to lower quality of mitochondrial gene annotation of the used genomes since contrastingly to the ortholog based results, we could annotate 36 out of 37 mitochondrial genes in our assembly.

The other gene losses concerned uncharacterized gene families as well as a ribonucleoprotein (IMP4), the claudin gene family encoding for tight junction proteins and a DNA polymerase.

We identified 12 gene gains and 84 losses reconstructed for the common ancestor of owls. We again found an expansion of the keratin gene repertoire (gain of one keratin and one scale-keratin like gene). GO parental terms of gains pointed again to metabolic changes but also those associated with the immune system. The gains contained also an olfactory receptor. The much more numerous losses were associated even at the higher level with many different GO categories again often related to metabolism (e.g., “regulation of amide metabolic process”, “pyridine−containing compound metabolic process”).

### Chromosome Scale Syntenies

Synteny with the chromosome-level assemblies of *Strix aluco* and *Bubo scandiacus* identified the Z chromosome of our male individual. It is the fifth largest chromosome in the Ural Owl assembly. The synteny between the two *Strix* genomes shows no major syntenic differences (Figure 7). This is also mostly true in the comparison to the Snowy Owl with the exception of the Z chromosome, which shows some internal rearrangements compared to the two *Strix* species. Whether this is caused by assembly quality and accuracy remains to be investigated. Compared to chicken and zebrafinch however, we identified several large-scale changes. We detected a fusion of chromosomes 5 and 6 of the Snowy Owl (corresponding to parts of chicken chromosome 4 and chromosome 5 and zebrafinch chromosome 4 and parts of 5, Figure 7 and Supplementary Table S6) into chromosome 4 of the two *Strix* assemblies. This is supported by previous cytogenetic analyses [27]. The remaining part of chicken chromosome 4 corresponds to Ural Owl chromosome 13, Snowy Owl chromosome 12 and zebrafinch 4a. The remaining part of chicken and zebrafinch chromosome 5 corresponds to two chromosomes in the Ural Owl (chromosomes 16 and 30). As in the zebrafinch, chicken chromosome 1 corresponds to two chromosomes in the Snowy Owl (chromosomes 2 and 6).

**Figure 7:**
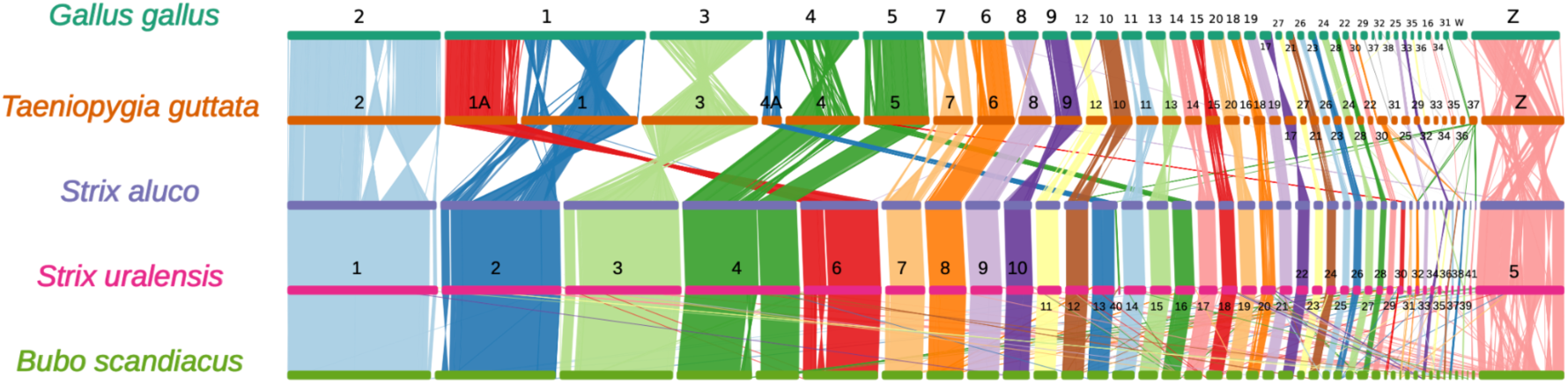
Chromosome scale synteny analysis. Synteny of chromosomes of *Taeniopygia guttata*, *Gallus gallus*, *Strix aluco* and *Bubo scandiacus* compared to the newly sequenced *Strix uralensis*. Syntenic regions amongst the species are indicated with a unique colour. Assignment of sex chromosomes was based on the *S. aluco* genome annotation. The chromosomes of all species have been reordered to highlight synteny relationships to *S. uralensis*. Plot made with NGenomeSynt.

For the remaining chromosomes of our Ural Owl assembly, we could mostly identify 1:1 relationships with chromosomes of Snowy Owl, chicken and zebrafinch with the exception of the Ural Owl microchromosomes 31, 35, 39 and 41 for which we could not unambiguously identify a corresponding chicken or zebrafinch chromosome but homology to Snowy Owl scaffolds and Tawny Owl chromosomes.

### Demographic history

The demographic history of the Ural Owl derived from our genome assembly appears to have a complex relationship to glacial and interglacial periods. The effective population was predicted to have decreased until around the Holstein interglacial period (3.74 × 10^5 – 4.24 × 10^5 ya) where its population size stabilized but remained low during the Saalian glacial period (4-1.3 × 10^5 ya) and began to increase as the Eemian interglacial period (1.3-1.15 × 10^5 ya) began to emerge. It continued to increase and reached a plateau during the last glacial period (Weichselian glaciation, 1.15-0.117 × 10^5 ya). Before the end of the last glacial period, at around 0.3 ×10^5 ya, the Ural Owl population began to decrease until it reached the current lowest effective population size (Figure 8).

**Figure 8.**
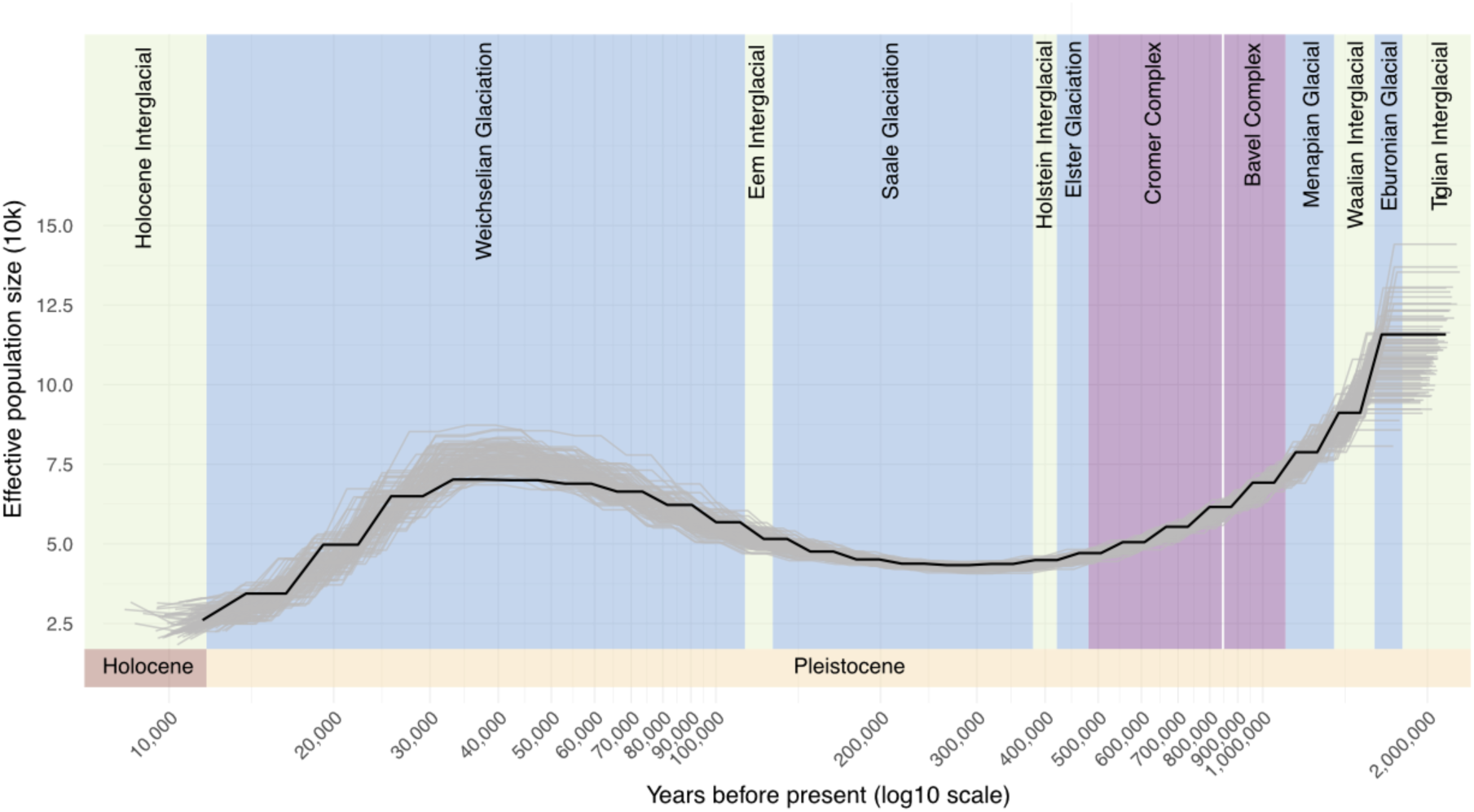
**Inferred demographic history of *Strix uralensis.*** The plot shows a Pairwise Sequentially Markovian Coalescent (PSMC) analysis based on the primary genome assembly. The x-axis shows years before present (ya) on a logarithmic scale and the y-axis shows the estimated effective population size. Bootstrap results are shown in light grey.

### Variation analysis over progressive cell passages

#### Karyotype confirms chromosome numbers and reveals no large variants caused by passaging

Chromosomal analyses detected 2n = 82 in both passages 5 and 10, corroborating the diploid chromosome number described for the Ural Owl previously (subspecies *S. uralensis uralensis* and *S. u. japonica*, [28]). No large-scale chromosomal rearrangements were observed between both passages (Figure 9).

**Figure 9:**
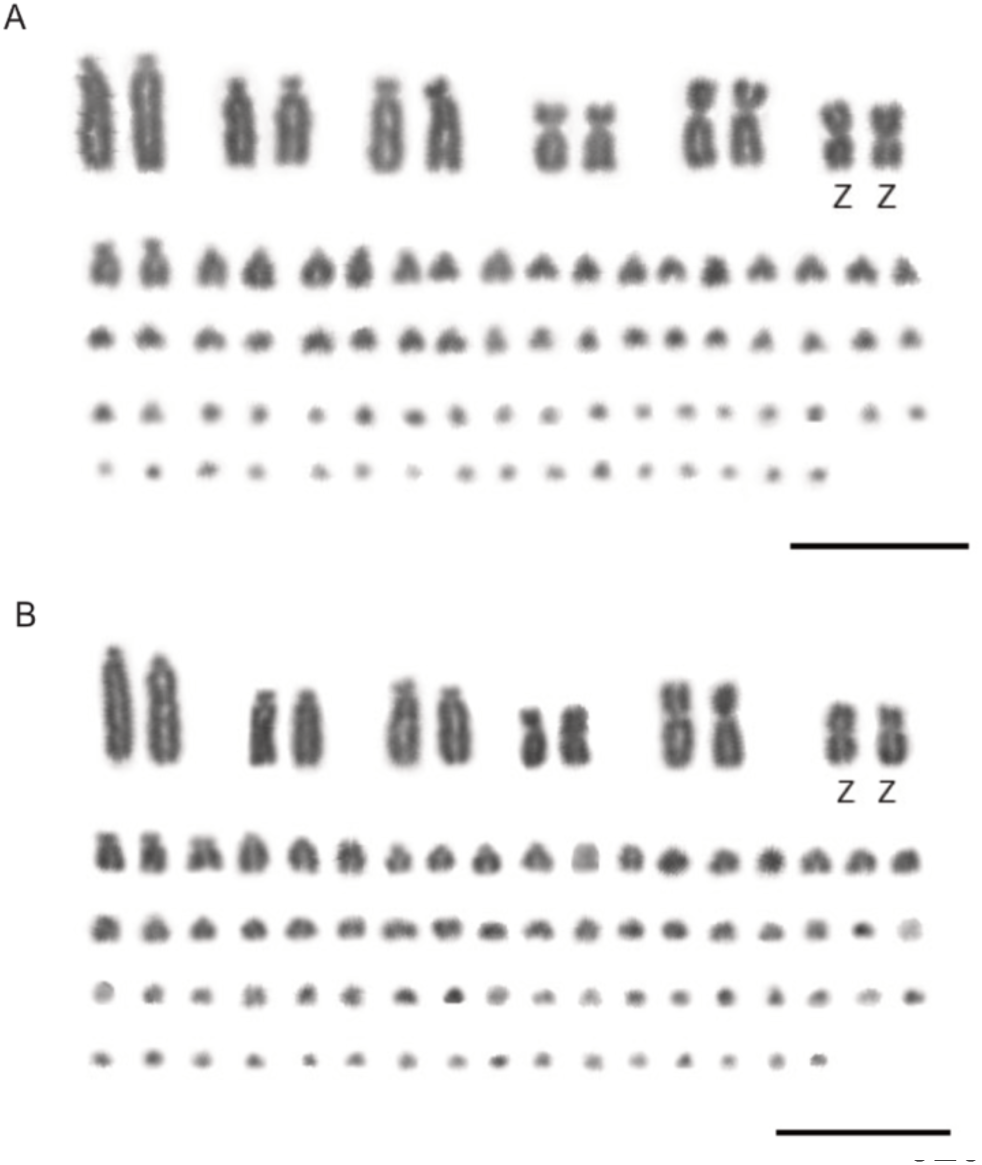
Karyotype analysis. Karyotype of *S. uralensis* male with 2n = 82 after passage 5 (a) and passage 10 (b). Bar = 10 μm.

#### Short-read variants

After quality filtering, we identified 885,159 variant sites (in the following referred to as SNPs) (Supplementary Figure S9). Out of these, the vast majority (i.e., 670,463 SNPs) were fixed variant sites across all samples and hence mostly represented heterozygous sites of the individual which are represented with just one of the two alleles in the reference genome or sites that had a wrong allele in the reference assembly.

The remaining 214,696 SNPs varied across samples, indicative of potential mutations, and were analysed in the following.

A comparison of all SNPs across all samples revealed that variant amount and type differed. The biggest differences resulted from SNPs called from the HiFi data as well as the passages 5.1 and 10.1 which appear to have more SNPs than the other passages. The majority of these are heterozygous first-alternate sites (Supplementary Figure S9, Table 2) and we suspect for many of those that they are false heterozygous calls rather than true mutations. To investigate this pattern further, we inspected genotype quality and depth focusing on sites with a genotype in a sample not found in any other sample (i.e., private sites) compared to the same metrics at all other sites of that individual (i.e., common variants and conserved heterozygous sites). This analysis revealed that all passages had similar median depth per variant site (DP ∼26.78 ± 6.55) and genotype quality (GQ ∼99), suggesting rather consistent data quality across samples (Table 2). It further showed that median depth and quality of private SNPs consistently had a significantly lower depth and quality than the average non-private site, suggesting that these alleles are to some extent wrongly called (Supplementary Figure S10). To account for patterns potentially driven by sequencing technology, we also assessed in each replicate at how many positions it differed compared to passage 0. This revealed the same pattern of an increase of SNPs in samples passage 5.1 and 10.1

**Table 2:** SNP statistics over progressive cell culture passages.

| Sample | Median depth of variant sites | Median GQ of variant sites | SNPs other than shared heterozygous sites/private to individual | SNPs other than shared heterozygous sites as a percentage of the total genome size [%] | SNPs that are 0/1 | SNPs that are not 0/1 nor 1/1 | SNPs other than fixed heterozygous sites [%] | SNPs different from passage 0 | SNPs different from passage 0 [%] |
| --- | --- | --- | --- | --- | --- | --- | --- | --- | --- |
| HiFi Reads | 39 | 99 | 78,983/35,645 | 0.0057 | 77,682 | 1,265 | 0.14 | - | - |
| Passage 0 | 27 | 99 | 57,929/8,212 | 0.0042 | 56,232 | 816 | 0.922 | - | - |
| Passage 5.1 | 27 | 99 | 78,193/23,889 | 0.0057 | 76,709 | 716 | 0.809 | 58,620 | 6.62 |
| Passage 5.2 | 29 | 99 | 55,225/3,280 | 0.0040 | 54,037 | 615 | 0.695 | 38,765 | 4.38 |
| Passage 5.3 | 28 | 99 | 57,664/5,581 | 0.0042 | 56,267 | 781 | 0.882 | 41,301 | 4.67 |
| Passage 10.1 | 20 | 99 | 103,086/63,058 | 0.0075 | 101,450 | 25 | 0.028 | 96,768 | 10.93 |
| Passage 10.2 | 28 | 99 | 54,404/3,410 | 0.0039 | 53,373 | 420 | 0.475 | 38,921 | 4.40 |
| Passage 10.3 | 28 | 99 | 55,976/3,758 | 0.0041 | 54,679 | 642 | 0.725 | 39,843 | 4.50 |
| Passage 10.4 | 15 | 99 | 54,215/15,521 | 0.0039 | 52,153 | 750 | 0.847 | 50,033 | 5.65 |

## Discussion

Reference genomes are accumulating across the tree of life and here birds have seen special attention fuelled by initiatives such as B10K (https://b10k.com/) [29,30]. Still, many of these genomes remain incomplete in terms of chromosomal-scale assembly type as well as gene annotation comprehensiveness. Genome assembly quality can impact phylogenomic inferences, analyses of gene prediction, gene family expansion and contraction, and most importantly structural evolution. In an effort to allow such analyses for the vastly understudied bird order Strigiformes, we here present a reference genome for the Ural Owl, that is among the best bird genome assemblies currently, reflected by assessments of sequence and gene completeness. We could place most of the genome into chromosomal-scale scaffolds, which are in line with the species karyotype, that we also confirm by cytogenetics. We further located the supposedly *Strix*-specific chromosomal fusion which distinguishes it from the genus *Bubo* [4]. Besides this, we identified several other chromosomal rearrangements between owl genomes (which are overall very syntenic) and those of chicken and zebrafinch. For four microchromosomes we could not identify a reliable homologous chromosome in chicken nor zebrafinch yet syntenic scaffolds in the Snowy Owl and microchromosomes in the Tawny Owl assemblies. In addition, the microchromosomes we assembled were also well supported by our HiC data. We thus hypothesize that the lack of synteny to chicken and zebrafinch might reflect more structural changes specific to owls. Matchingly, a first analysis of the Ural Owl genome content also indicates an important increase in repetitive sequences compared to most other non-owl bird genomes. Birds on average have rather compact genomes compared to other vertebrate lineages (average ∼1.1 Gb [31]), mostly owing to a low content of repetitive elements (10-15 %). Until now, owls were seemingly no exception to this with an average genome size of 1.2 Gb and a repeat content of ∼8,6 % [31]. Still, cytogenetic studies already suggested differently and hint at owls being rather an exception in the avian lineage with large scale variations in karyotypes, which our assembly also supports. For example, the barn owls have chromosomes of a more homogenous smaller size while the true owls have a more classical avian karyotype with macro-and microchromosomes, suggesting chromosome fusion and fission in owls [4]. Interestingly, the recently published genome (1.6 Gb) of the Snowy Owl further extends this observation by demonstrating that it has one of the highest reported repeat contents for birds (28.34 %), mainly composed of centromeric satellite DNA [32]. Our assembly’s total repeat content, at 21.07 % (1.5 % of which is unidentified), follows this pattern. While both owl genomes’ repeat expansions are largely driven by retrotransposons, the Snowy Owl had a stronger increase of LTRs compared to LINEs than the Ural Owl. Nevertheless, LTR retrotransposons are the largest repeat class also in the Ural Owl, and, especially the youngest repeat expansion is also driven by LTRs suggesting this pattern to be more broadly present in true owls. In the Snowy Owl, the repeats are suspected to have driven the evolution of novel centromeres. Accordingly, cytogenetic analyses already identified large centromeric satellite blocks shared among and unique to true owls [27]. Other bird lineages with increased repeat content are woodpeckers and the Common Scimitarbill (*Rhinopomastus cyanomelas*) [31]. The cause and consequences of the repeat extensions in the genera *Strix* and *Bubo* remain unclear at this point, which is also true for the woodpecker [33].

The number of microchromosomes identified in *S. uralensis* (n = 31) is consistent with the average number of microchromosomes reported by Tegelström and Ryttman [34] from karyotypes of over 230 bird species. However, there is no established rule for distinguishing between macrochromosomes and microchromosomes (e.g. [35]). We here confirm that in birds, microchromosomes have a higher gene density than macrochromosomes (e.g. [5,35–37]). We could further show that the higher density of genes on microchromosomes is associated with a reduction of gene length, which in turn is due to correspondingly shorter introns. A similar pattern has been reported in chickens, where the size of the chromosome correlates with the length of the genes it harbours [38]. Thus, in true owls, microchromosomes hold up the crucial role they supposedly have played throughout vertebrate evolution [39].

Our assembly seems to be particularly well-suited for an analysis of gene content due to a high completeness of gene annotation. However, due to vastly varying assembly qualities, the correctness of our gene family expansion analysis should be taken with a grain of caution. Still, this preliminary analysis suggests that in the future we will be able to connect changes in gene content to adaptations of owls. This is supported by Ural Owl specific gene gains in the GO-term derived function of e.g. “eye development”, “sensory organ development” and “neurogenesis”, which could be linked to adaptations required for a nocturnal, predatory lifestyle [40].

We also offer candidate genes for further investigation that characterize predatory lifestyle, i.e., genes gained in the common ancestor of Strigiformes and Accipitriformes that acquired this lifestyle. We especially observed gains of genes with a metabolic function which could relate to the change in diet in the ancestor of these two bird orders. We also found gains of keratin genes, in the common ancestor of the two predatory bird lineages and also in the ancestor of owls. Feathers are epidermal appendages. Vertebrate skin appendages consist of two fibrous proteins, alpha and beta keratins. Interestingly, β-keratins are exclusively found in reptiles and birds. Both keratin gene families show expansions in different lineages. The Barn Owl had the lowest number (6) of β-keratins in a study comparing 48 bird (draft) genomes. The zebra finch in comparison had 149 genes [41]. This comparison further showed that the proportion of claw β-keratins and keratinocyte β-keratins is higher in predatory birds. We support the latter finding with the detection of three gains of feather and scale keratins in the common ancestor of Accipitriformes and Strigiformes and two further gains in the ancestor of all owls. These keratin genes deserve more attention as potential candidates that could underlie morphological adaptations of feathers in predatory birds in general but also more specifically in the mostly nocturnally hunting owls. Their silent flight is made possible by physical characteristic fringes of the feathers on the leading edge of the wings [42]. The genomic basis of this adaptation remains to be identified.

The Ural Owl is protected under the CITES convention Annex II and the Bern Convention on the Conservation of European Wildlife and Natural Habitats. While globally not (yet) under concern, the species went extinct in Germany and Austria due to habitat destruction but also direct persecution. Reintroduction programmes have been started, however due to low availability of breeding couples, individuals of various origins are used for these actions [18]. An analysis of marker genes neither supported morphological subspecies nor did it reveal a phylogeographic population structure for the Ural Owl; yet, it revealed genetic clusters that could be informative for breeding programs [12]. The here generated reference genome will facilitate future genomic studies in this direction of *S. uralensis*.

With an estimated genome wide heterozygosity of 0.2 % (2 het/kb), the here sequenced individual shows a higher level of heterozygosity than genomes of endangered bird species (red list status accessed March 2025, /www.iucnredlist.org/) such as the white-eared night heron (*Gorsachius magnificus*, Endangered, 0.49 het/kb) [43], Andean condor (*Vultur gryphus*; Vulnerable, 0.75 het/kb) and California condor (*Gymnogyps californianus*; Critically Endangered, 1.34 het/kb) [44], and Crested ibis (*Nipponia nippon*, Endangered, 0.043 het/kb) [45]. A similar heterozygosity level as the one we estimated for the Ural Owl was detected in e.g., Wild Turkey (0.24 %) and Mallard (0.26 %) [46]. It is somewhat lower than levels reported for other Strigiformes such as little owl (*Athene noctua*; 0.593) [47], Tawny Owl (*S. aluco*, 0.57 to 0.70) [48] and Barn Owl (*Tyto alba*, 0.59 to 0.71) [49] yet twice as high as in the Burrowing Owl (*Athene cunicularia*; 0.1%) [50].

Overall, species with a threat of going extinct show reduced levels of heterozygosity compared to non-threatened related taxa [51]. Related taxa of the same bird order with and without risk of extinction, differ quite drastically in genome-wide heterozygosity [45]. These differences likely result in lower evolutionary potential, reduced reproductive fitness and may contribute to species extinction [51]. It remains to be assessed at which level heterozygosity reduction causes an issue for a particular species. By generating genomic data for the Ural Owl, we contribute to the required knowledge for genetic monitoring of biodiversity.

Further, our reference genome already sheds light on the demographic history of the species, indicating both population contractions and expansions, apparently related to ecological effects of the glacial-interglacial cycle. In particular, the pattern over the last 120,000 years not only demonstrates the Ural Owl’s tolerance to lower temperatures, but more importantly reflects its flexible habitat choice of semi-open woodlands with a mixed composition of broadleaf and coniferous species [10]. From the Eemian interglacial through the Weichselian glaciation, climatic changes caused fluctuations in ice sheet extent and associated changes in vegetation composition, including a gradual and/or repeated reduction in forest cover leading to a treeless shrubby or grassy tundra from the mid-Weichselian (e.g., [52–54]). While the open or semi-open structure of the woodland habitats favoured the Ural Owl’s preference for breeding and hunting grounds [11] until the mid-Weichselian, the expanding tundra substantially reduced suitable habitats, leading to a marked decrease in effective population size.

In the light of species preservation, protection and restoration, *ex situ* efforts are gaining more attention. Cell culturing is a valuable and widely used technique, spanning applications from basic science to biotechnology research [25]. However, there is no consensus regarding the number of passages considered “safe” before cells experience metabolic changes, DNA damage, and chromosomal instability. What is deemed “high passage” for one cell culture may not lead to significant passage effects in another [26]. Thus, the effects of prolonged culture are complex and depend on the individual cell culture, tissue, and species.

The first criterion for identifying healthy and stable cells is observing cell morphology. Chromosome content serves as another critical benchmark, as normal cells maintain a stable chromosome number. Some studies using non-model species, such as felines [55] and fishes [56], showed no heteroploidy on karyotypes obtained by cell cultures. However, to the best of our knowledge, this is the first study that addresses genomic and chromosome changes in wild birds and compares the effect of different cell passages on genome integrity.

Cryopreservation of cells has increasingly been considered a strategy for conservation as new technologies using genetic material from somatic cells (e.g., somatic cell nuclear transfer or induced pluripotent stem cells) are evolving [23,57]. One of the prerequisites for nuclear donor cells and *ex situ* conservation is the stability of chromosomes [55]. Studies that investigate cell line passage and age effects are still scarce in non-model organisms and are crucial since altered metabolism and genomic instability no longer represent reliable models of their original source of material.

Herein, comparison between karyotypes of passage 5 and passage 10 showed no differences, suggesting that no large structural rearrangement occurred during the progressive number of passages and that it is safe to establish the diploid number of (this) bird species until at least the 10th passage. Genetic instability is well-documented in cells that have undergone more than 20 passages, particularly in transformed continuous cell lines (e.g., [58]) and tumour cell lines [59,60]. However, for primary cell cultures, a straightforward method to determine the safest passage number before cells develop mutations or genetic instability is lacking. We opted to cultivate cells up to passage 10 based on two factors: first, the uncertainty surrounding the exact passage limit at which primary cells may enter senescence (as non-continuous cell line has a limited *in vitro* life time); and second, technical challenges observed during later passages as cells began to exhibit signs of morphological decline, including the presence of granules and debris, difficulty detaching, and a reduced growth rate, all of which would complicate further subculturing beyond passage 10. This seems to suggest rather safe cell culturing for this species until passage 10. To some extent, this is supported by our SNP analysis of several passage replicates which indicated no general pattern for increased genomic changes between passages 5 and 10. However, we detected outlier samples with respect to SNP numbers among replicates of both passage numbers. At this point, we lack any point of reference expectation as to how many (potential) mutations are to be expected in a cell culture system as the one we applied. Compared to overall levels of variant sites, the number of SNPs in the individual samples which could be mutations is lower and represented between 0.0039-0.0075 % of the genome assembly length. The effects of these variants remain to be determined as well as the reason for between replicate differences. We further suspect that several mutations are variant-calling artefacts, supported by lower SNP calling quality, which asks for an exploration of mutation identification and more importantly validation for the type of cell culture we have set up here.

### Potential implications

We were able to assemble a reference genome for the Ural Owl of gold standard quality which is open to the community to be used for broader comparative genomic studies and phylogenomic analysis but also serves immediately to researchers interested in the Ural Owl for taxonomic and conservation aspects. With the data generated, we contribute to the endeavour of sequencing all life on Earth https://www.earthbiogenome.org/. Our analysis of genomic data derived from cell passages opens space for discussion of cell cultures as material for genomics especially for species with limited biological material available. The workflows applied by us could be used on similar data from other species.

## Methods

### Species origin and sampling strategy

Skin and muscle tissue samples from a ten-year-old male individual of *S. uralensis* (ring ID ZG-14.0-10-0234) were obtained from the Raptor Center & Wildlife Park Hellenthal (Wildfreigehege und Greifvogelstation Hellenthal, Hellenthal, Germany) during necropsy in 2020. The procedure was performed by Dominik Fischer, who is a veterinarian and approved to handle animals. No further approval was needed for this study. DNA barcoding was performed (collection ID ZFMK-TIS-50475) to ensure species identity using primers for COI from Astrin and Stüben [61] and sequences matched against BOLD (Barcode of Life Data System) [62]. The barcode sequence has been uploaded to BOLD as FOGS049-22.

### Reference genome

#### Sequencing

DNA was extracted from the skin biopsy (collection ID ZFMK-TIS-50482, stored at LIB Biobank in liquid nitrogen vapor phase) using the Monarch HMW DNA Extraction Kit (NEB, Ipswich, USA). High-molecular weight status was validated by quality control with capillary electrophoresis (Agilent Femto Pulse) and a SPK 3.0 PacBio HiFi library was prepared according to the recommendations by the vendor. Next, HiFi SMRT sequencing was performed on two SMRT cells on a PacBio Sequel IIe (Pacific Biosciences, Menlo Park, USA) at the Max-Planck Genome-centre Cologne (MP-GC; Cologne, Germany). Also, a chromatin-capture library was prepared from cryopreserved cells generated for the analysis over progressive cell passages as described below with an Arima-Hi-C Kit according to the protocol for Mammalian Cell Lines followed by sequencing on an Illumina NextSeq 2000 in paired-end read mode.

### Read processing

#### HiFi data

Contaminant sequences were filtered from the HiFi reads using Kraken2 v2.1.3 [63,64] with the Kraken database kraken2 PlusPFP downloaded in March 2023 and parameters “--confidence 0.51--use-names”. HiFi read quality was assessed using seqkit v2.8.2 [65,66] and a k-mer-based approach. K-mers were calculated with Meryl v1.4.1 [67] using the parameters “count k=21”, and the counts were converted into a histogram with the *meryl histogram* command.

To verify the ploidy of the individual, Smudgeplot v0.2.5 [68] was used. First, k-mers within a specific range (lower-upper), determined with the *smudgeplot.py* cutoff function, were extracted using the meryl *print less-than* command. These filtered k-mers were then processed with *smudgeplot.py hetkmers* to calculate the coverage of unique heterozygous k-mer pairs. The resulting coverage was plotted using smudgeplot_plot.R.

GenomeScope2 v2.0.1 [68] with default parameters was used to estimate genome size, heterozygosity, and the homozygous and heterozygous coverage peaks.

ROHan v1.0.1 [69] was used to identify large (>1 Mb) runs of homozygosity.

#### Hi-C data

Adapter removal and quality filtering of the raw Hi-C reads were performed using Fastp v0.23 [70] with parameters “--length_required 95,--qualified_quality_phred 20--adapter_fasta”, with a curated adapter list of the most common adapters used as input.

Error correction was done using *Tadpole* from BBMap v39.01 (https://sourceforge.net/projects/bbmap/), with parameters “k=50, reassemble=t, mode=correct, minprob=0.6, prefilter=1, prehashes=2, and prealloc=t”. To remove contamination from the short, Hi-C reads, Kraken2 v2.1.3 was used similarly to the HiFi reads but in paired-read mode, with parameter “--paired”.

#### Initial Genome Assembly

The HiFi reads were used with Hifiasm v0.19.5 [71] to generate a phased genome assembly. In order to obtain an optimal, phased genome we tested several Hifiasm parameters before choosing the ones that provided us with the assembly of the highest contiguity and completeness with both phased haplotypes having a similar length. We tested all possible combination of different purging level (0, 2, 3), increasing run-time, and number of iterations (“--n-weight 5--n-perturb 50000--f-perturb 0.5-D 10-N 150-s 0.2”) and explicitly providing the homozygous peak to Hifiasm which was estimated by GenomeScope (“--hom-cov 40”) (for more details see Supplementary Table S1).

#### Genome Scaffolding

The selected haplomes from Hifiasm were split at positions containing Ns using *split_fa* from the Purge_Dups package v1.2.6 [72]. The resulting sequences were mapped to themselves using Minimap2 v2.26 [73] with parameters “-x asm5-DP” and to the HiFi reads using Minimap2 with parameters “-x map-hifi”. These mappings were used to remove assembly duplicates with Purge_Dups.

#### Mitochondrial Genome Detection

To identify and extract the mitochondrial genome, we utilized MitoHiFi v3.2.1 [74,75] referencing the sequence NC_038218.1 from *S. uralensis* (isolate C5 mitochondrial genome, complete, https://www.ncbi.nlm.nih.gov/nuccore/NC_038218.1). The most likely scaffold was kept and identified as the mitochondrial chromosome (MT) and all other candidate scaffolds were removed from the assembly.

#### Assembly Manual Curation

Hi-C reads were aligned to the final assemblies and a Hi-C contact map was created using PretextMap v0.0.2 (https://github.com/sanger-tol/PretextMap). A HiFi coverage track was generated from the aligned HiFi reads using bedtools *genomecov* and integrated into the Pretext map with *PretextGraph*.

Manual curation was performed within PretextView v0.0.2 (https://github.com/sanger-tol/PretextView), where scaffolds were reordered and oriented based on Hi-C interaction frequencies. Following curation, the final assembly scaffolds were processed with AGP tools from the Vertebrate Genomes Project (VGP) using the rapid manual curation protocol (https://gitlab.com/wtsi-grit/rapid-curation/-/tree/main) established by the Darwin Tree of Life consortium (https://www.darwintreeoflife.org/) to create the curated assembly. Scaffold names were further sorted and renamed by size using a combination of seqkit v2.8.2 and SAMtools v1.19.2 [76]. The final Hi-C contact map was visualized with HiCExplorer [77].

#### Genome Quality Control

The completeness of the final curated assembly was assessed using BUSCO v5.8 [78,79]] and compleasm v0.2.6 [80] with the aves_odb10 lineage. Assembly contiguity and general assembly metrics were calculated using Quast v5.2.0 [81].

For k-mer-based analysis, k-mer counts were generated for each assembly using Meryl. These counts were analysed with Merqury v1.3 [82] to estimate assembly completeness and accuracy. The analysis yields Merqury’s consensus QV, which is estimated by comparing the read and assembly k-mer counts and then transformed to a log-scaled probability of base-call errors. A higher QV indicates a more accurate assembly. We also obtained a Merqury completeness percent, which reflects the proportions of high-quality HiFi read k-mers present in the assembly.

HiFi reads were mapped to each assembly using Minimap2 with parameters “-ax map-hifi”. Alignment quality and coverage distribution were assessed using Qualimap v2.3 [83].

Potential contamination and quality was also assessed using the blobtoolkit pipeline v3.5.4 [84] and visualized using the interactive Blobtoolkit viewer in the Galaxy EU server [85].

### Genome Annotation

#### Repeat annotation

Repetitive elements in the primary assembly were identified and annotated using EarlGrey v5.1.1 [86], which was run with RepeatMasker v4.1.5 [87] and RepeatModeler v2.0.6 [88]. In addition to the RepeatModeler library, we used a previously-created, custom avian TE library to mask repetitive elements [89]. The softmasked genome was used for protein-coding gene prediction.

#### Protein-Coding Gene Annotation

To perform protein annotation, we created two reference protein sets. Set one contained only the merged proteomes of the following publicly available genomes, downloaded using the NCBI dataset cli v16.3.0 [90]*: S. nigrolineata* (GCA_013396715.1), *Gallus gallus* (GCF_016699485.2), *Glaucidium brasilianum* (GCA_013399595.1), *Falco peregrinus* (GCF_023634155.1), *Athene cunicularia* (GCF_003259725.1), *Aquila chrysaetos* (GCF_900496995.4) [91], *Taeniopygia guttata* (GCF_003957565.2), and *Anas platyrhynchos* (GCF_015476345.1).

Protein set two was created by merging all proteomes in set one with proteins from the following public and curated databases: i) proteins from the BUSCO v5.4 aves_odb10 dataset, ii) aves proteins from OrthoDB v11 [92] were obtained using Tomas Bruna’s orthodb-clades pipeline (https://github.com/tomasbruna/orthodb-clades) and iii) proteomes were also extracted from the UniProt database for the following species: *Calypte anna* (UP000054308), *Steatornis caripensis* (UP000516988), *Cnemophilus loriae* (UP000517678), *Dasyornis broadbenti* (UP000521322), *Corythaixoides concolor* (UP000526942), *Irena cyanogastra* (UP000530962), *Bucco capensis* (UP000534107), *Cephalopterus ornatus* (UP000543364), *Molothrus ater* (UP000553862), *Ptilonorhynchus violaceus* (UP000584880), *Promerops cafer* (UP000587587), *Vidua chalybeata* (UP000634236), and *Urocolius indicus* (UP000654395).

Protein-coding genes in the *S. uralensis* genome were annotated using a combination of *ab initio*, protein similarity, and transcriptome-based protein prediction models. BRAKER3 v3.0.3 [93,94] was run in EP mode using the protein set two described above. GALBA v1.0.11.2 [95] was run using protein set one to annotate genes. The outputs from GALBA and BRAKER3 v3.0.2 were combined using *TSEBRA* from the BRAKER3 package. To ensure high-quality annotations, only the longest gene orthologs for each locus were retained using the *agat_sp_keep_longest_isoform.pl* script from the AGAT package v1.4.1 [96].

### Demographic History of *S. uralensis*

The demographic history of the Ural Owl was reconstructed using PSMC v0.6.5 as implemented by [97]. Variants were called per chromosome using a combination of BCFtools v1.21 (http://github.com/samtools/bcftools) *mpileup* with parameters “-Q 30-q 30” and bcftools call using the “-c” option. The resulting VCF file was converted to a consensus fastq format using the *vcfutils.pl* vcf2fq script with parameters “-d 10,-D 60, and-Q 30”. The PSMC model was run with the following parameters:-N25-t15-r5-p “2+2+25*2+4+6” and 100 bootstraps, a generation time of 3 years [98,99] and an assumed mutation rate of 4.6×10-9 [100,101].

### Genome Synteny

In order to identify the Z sex chromosome within our genome assembly and to assess the synteny of different bird genomes we used the *GetTwoGenomeSyn.pl* built-in script of NGenomeSyn [102] with options: “-MappingBin minimap2-MinLenA 100000-MinLenB 100000-NumThreads 5-MappingPara ‘-Lx asm10--eqx-I 200G--MD-N 1’” to estimate chromosome-scale alignments between *G. gallus* (GCA_024206055.2)*, T. guttata* (GCF_048771995.1), *S. aluco* (GCA_031877795.1), *S. uralensis,* and *B. scandiacus* (GCA_965212795.1). To improve visualisation and reduce noisiness in the synteny plot, we used the get.synteny.blocks.multi function of the asyntr R package (https://github.com/simonhmartin/asynt) to aggregate single alignments into synteny blocks of incrementally increasing minimum block size of 100 to 4000 and filtered for chromosomes smaller than 1000 bp and for alignment lengths smaller than 200 bp. For visualization purposes, we manually removed alignments between *G. gallus* and *T. guttata* that were not supported by Luo et al. [103].

### Functional Gene Annotation

Predicted genes were functionally annotated by performing sequence similarity searches against the Swiss-Prot database using *BLASTP* from BLAST v2.13.0+ with default parameters. As with our own genome’s annotation, we used *agat_sp_keep_longest_isoform.pl* to only keep the longest isoform of each gene locus from the *Aquila chrysaetos, Gallus gallus, S. nigrolineata*, *Athene cunicularia, Glaucidium brasilianum, Falco peregrinus,* proteomes of protein set one together with *Tyto alba* (GCA_018691265.1) and *Haliaeetus leucocephalus* (GCA_000737465.1) and used OrthoFinder v2.5.5 [104] to estimate orthologous gene families among those species. This analysis identified orthogroups and genes that have undergone expansion or contraction in the Ural Owl, as well as orthogroups unique to this species.

### Gene Ontology Term Analysis

Gene Ontology Term analysis was performed by mapping all Ural Owl genes to the Vertebrate Eggnog database using eggnog mapper v2.1.12 [105,106]. Missing GO Terms were filled in with the GO Terms of the previously found Swiss-Prot gene symbols associated with each gene. Next, the genes belonging to gene families unique to the Ural Owl (found with Orthofinder) were analysed by Revigo v1.8.1 [107] with default settings and choosing the *Large* subset option. The resulting GO Terms were analysed with Revigo again with default settings and this time with the *Small* subset option. The full *Biological Process* Revigo table was plotted in R v4.4.2 using an edited version of the Revigo treemap plotting script.

### Variation analysis over progressive cell passages

#### Cell Culture

Primary cells were grown from a skin biopsy of the same individual as used for genome sequencing (collection ID ZFMK-TIS-51054) previously stored at LIB Biobank in liquid nitrogen, following standard protocols. Skin tissues were rapidly thawed, minced into small fragments, and transferred to cell culture flasks. Flasks were incubated at 37°C with 5 % CO_2_ in Fibroblast Growth Basal Medium (FBM; Lonza, Cologne, Germany) supplemented with 20% Fetal Bovine Serum (FBS; Biowest, Nuaillé, France) including antibiotics (100 U/mL penicillin and 100 g/mL streptomycin; Sigma-Aldrich, St. Louis, United States). Cells were visually inspected in inverted microscope Nikon Eclipse TS2 for contamination and cell media was changed every 2-3 days. After reaching ∼80 % confluence (determined visually), cells were propagated using 0.125 % trypsin solution (Biowest), at subculture ratio 50:50. Cells were harvested for DNA extraction and chromosome analysis at passages 5 (three different replicates) and passage 10 (four different replicates).

#### Chromosome sampling for large variant analysis

In order to investigate the stability of the karyotype composition through different passages, chromosome preparations were obtained from cells for passages 5 and 10, according to [108], with modifications. Chromosomes were harvested after treatment with colchicine 0.01% for one hour, followed by hypotonic treatment with 0.075 M KCl, and cell fixation in methanol / acetic acid (3:1). Slides were stained with Giemsa 5 %. At least 20 metaphases for each passage were analysed to define the diploid number (2n) in a Zeiss microscope Axio Imager Z2m.

#### DNA extraction of primary tissue and cell culture passages

Passage samples were extracted using the DNeasy Blood & Tissue Kit (Qiagen, Hilden, Germany) following the manufacturer’s protocol for cultured cells, while muscle tissue of the same individual (collection ID ZFMK-TIS-50476) previously stored at LIB Biobank in 96 % ethanol (passage 0) was extracted using the standard protocol of the same kit.

#### Sequencing of primary tissue and passages

After DNA extraction, samples were sent for purification (Vahtstm DNA Clean Beads; Vazyme Biotech, Nanjing, China), PCR-free library preparation (NEBNext Ultra II FS DNA PCR-free Library Prep Kit for Illumina; NEB) and subsequent paired-end sequencing on a NovaSeq 6000 (Illumina, San Diego, USA) using the NovaSeq 6000 S4 Reagent Kit (Illumina) to Biomarker Technologies (bmkgene; Beijing, China).

#### Read Mapping

The Illumina reads of all passages (passage 0, i.e. the primary tissue, three passage 5 replicate samples and four passage 10 replicate samples) were processed with fastp v0.20.0 [70] with parameters “--length_required 95,--qualified_quality_phred 20--adapter_fasta”, with a curated adapter list of the most common adapters used as input and decontaminated with Kraken2 v2.1.3 with the Kraken database kraken2 PlusPFP database downloaded in March 2023 in paired-read mode, with parameter “--paired--confidence 0.51--use-names”. These reads were then mapped to the reference genome using BWA-MEM2 v2.2.1 [109,110] with the *mem* command and options “-M-R”, where a read group (RG) specific to each sample was used for “-R”. The resulting output was sorted using SAMtools v1.19.2, and additional processing steps (SAMtools’ *fixmate*, *sort*, and *markdup*) were performed to generate the final mapping files for each sample.

#### SNP Calling

SNP calling for each cell passage sample and the HiFi reads (“reference”) was performed individually using GATK HaplotypeCaller v4.2.6.1 [111,112] with the options “-ERC GVCF--min-base-quality-score 30--pcr-indel-model NONE”. Joint SNP calling was performed by first combining samples using GATK *GenomicsDBImport* with the option “--batch-size 3”. The combined database was then used for joint SNP calling with GATK’s *GenotypeGVCFs*.

Variant quality recalibration was conducted in three rounds. GATK’s *BaseRecalibrator* was run with the option “--maximum-cycle-value 50000”, followed by GATK’s *ApplyBQSR* for each sample before re-calling variants individually and collectively. From the final set of called genotypes, SNPs were extracted using GATK’s *SelectVariants* with the option “-select-type SNP” and filtered with GATK VariantFiltration using the filters: “QD < 2.0, FS > 60.0, MQ < 40.0, SOR > 3.0, MQRankSum <-12.5, ReadPosRankSum <-8.0, QUAL < 30.0”.

Variants were filtered for depth, minor allele frequency (MAF) and the fraction of missing genotypes using BCFtools filter v1.21 (https://github.com/samtools/bcftools) with the options “-e”INFO/DP<$MIN_DEPTH || INFO/DP>$MAX_DEPTH“” and“-i”MAF>$MAF && F_MISSING<=$MISS“.

#### Short-read variant analysis in passages

To assess the quality of the DNA contained in cell cultures and to understand its potential for being used as an amplified genomic resource, these filtered SNPs were then analysed in R v4.4.2. Passages 5 and 10 were compared to passage 0 and the HiFi reads at sites where passages 5 and 10 differ from either of the reference passages. A Wilcoxon test from rstatix v0.7.2 was performed to test whether the depth and GQ of these SNPs of each sample were significantly different from the average DP or GQ. The variant calls were re-coded so that: “0|0” = 0, “0|1” = 1, “1|0” = 1, “1|1” = 2, “0|2” = 3, “2|0” = 3, “1|2” = 4, “2|1” = 4, “2|2” = 5, “0|3” = 6, “3|0” = 6 and plotted with ggplot2.

## Availability of source code and requirements

Project: Strix uralensis assembly, annotation and comparative analysis Location: Zenodo DOI: 10.5281/zenodo.15100180

Operating system(s): e.g. Platform independent Licence: CC0

## List of abbreviations

2n: diploid chromosome number
b: bases
bp: base pair
BLAST: basic local alignment search tool
BOLD: Barcode of Life Data System
BUSCO: Benchmarking Universal Single-Copy Orthologs
C: Celsius
CBD: convention on biological diversity
CITES: convention on international trade in endangered species of wild fauna and flora
DP: depth
EU: European Union
FBS: Fetal Bovine Serum
GATK: Genome Analysis Toolkit
Gb: gigabases
GO: gene ontology
GQ: genotype quality
Hi-C: high-throughput chromosome conformation capture
HiFi: high-fidelity
HMW: high molecular weight
INSDC: International Nucleotide Sequence Database Collaboration
Kb: kilobases
LINE: long interspersed nuclear element
LTR: long terminal repeat transposable element
M: molar
MAF: major allele frequency
Mb: megabases
MT: mitochondrial chromosome
PSMC: Pairwise Sequentially Markovian Coalescent
QV: quality value
RG: read group
ROH: runs of homozygosity
SNP: single nucleotide polymorphism
TE: transposable element
VCF: variant call format
VGP: Vertebrate Genomes Project
ya: years ago

## Data Availability

The sequencing reads, assembly and BioSample data supporting the results of this article are available in the INSDC under the BioProject number PRJNA1212906. Further datasets and code supporting the results of this article are available from Zenodo under DOI: <u>10.5281/zenodo.14676512</u>. Code is available from Zenodo under DOI: <u>10.5281/zenodo.15100180</u>.

## Declarations

The primary tissue used for this work was derived from a naturally deceased bird and provided by a veterinarian. We did not perform animal experimentation.

## Authors’ contributions

IC: conceptualization, data curation, formal analysis, investigation, methodology, software, validation, visualization, writing (original draft; review & editing); AM: conceptualization, data curation, formal analysis, investigation, methodology, validation, visualization, writing (original draft; review & editing); CBDN: conceptualization, formal analysis, investigation, methodology, writing (original draft; review & editing); DF: resources, writing (review & editing); NS: investigation, writing (review & editing); LvdM: investigation, writing (review & editing); BH: investigation, writing (review & editing); JJA: conceptualization, funding acquisition, supervision, writing (original draft; review & editing); TT: validation, supervision, writing (original draft; review & editing); AB: conceptualization, data curation, validation, visualization, funding acquisition, supervision, writing (original draft; review & editing).

## Acknowledgements

We thank Juliane Vehof and Benjamin Wipfler for enabling us to use their microscope.

