## Supplementary Figures S1-S10 for "A high-quality reference genome for the Ural Owl (*Strix uralensis*) enables investigations of cell cultures as a genomic resource for endangered species"

Content:

Supplementary Figure S1

Supplementary Figure S2

Supplementary Figure S3

Supplementary Figure S4

Supplementary Figure S5

Supplementary Figure S6

Supplementary Figure S7

Supplementary Figure S8

Supplementary Figure S9

Supplementary Figure S10

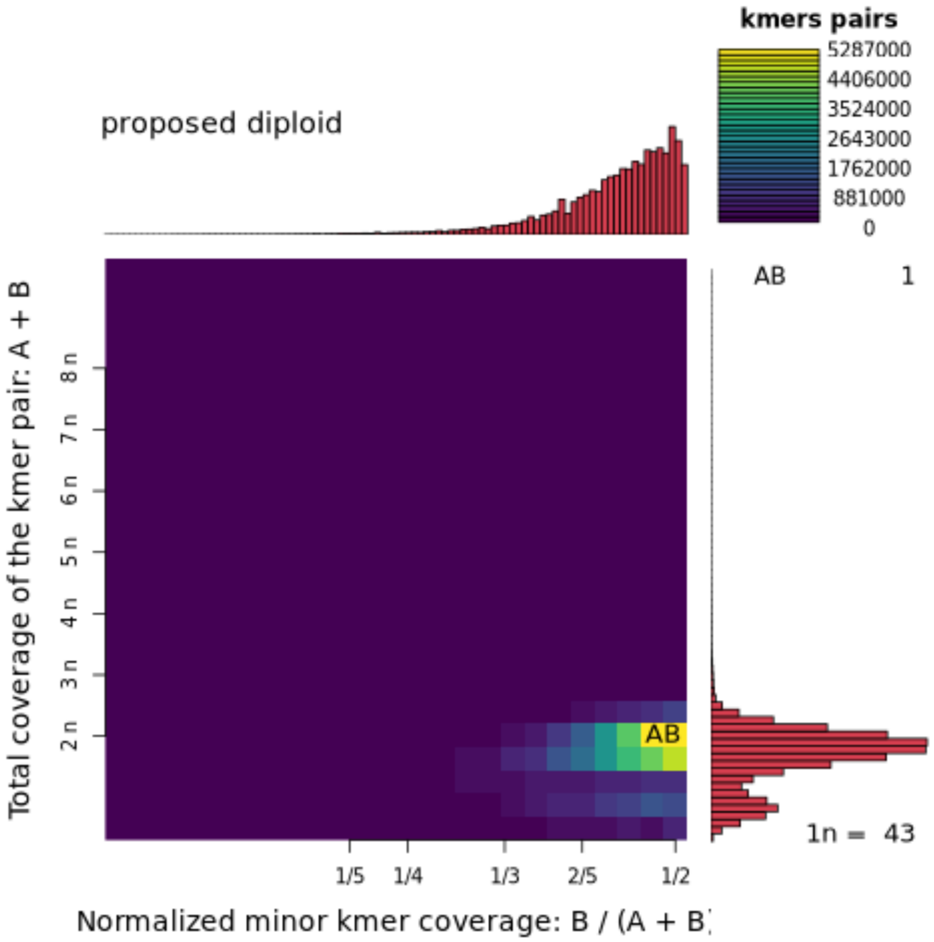


Supplementary Figure S1: PloidyPlot of *Strix uralensis* HiFi reads.

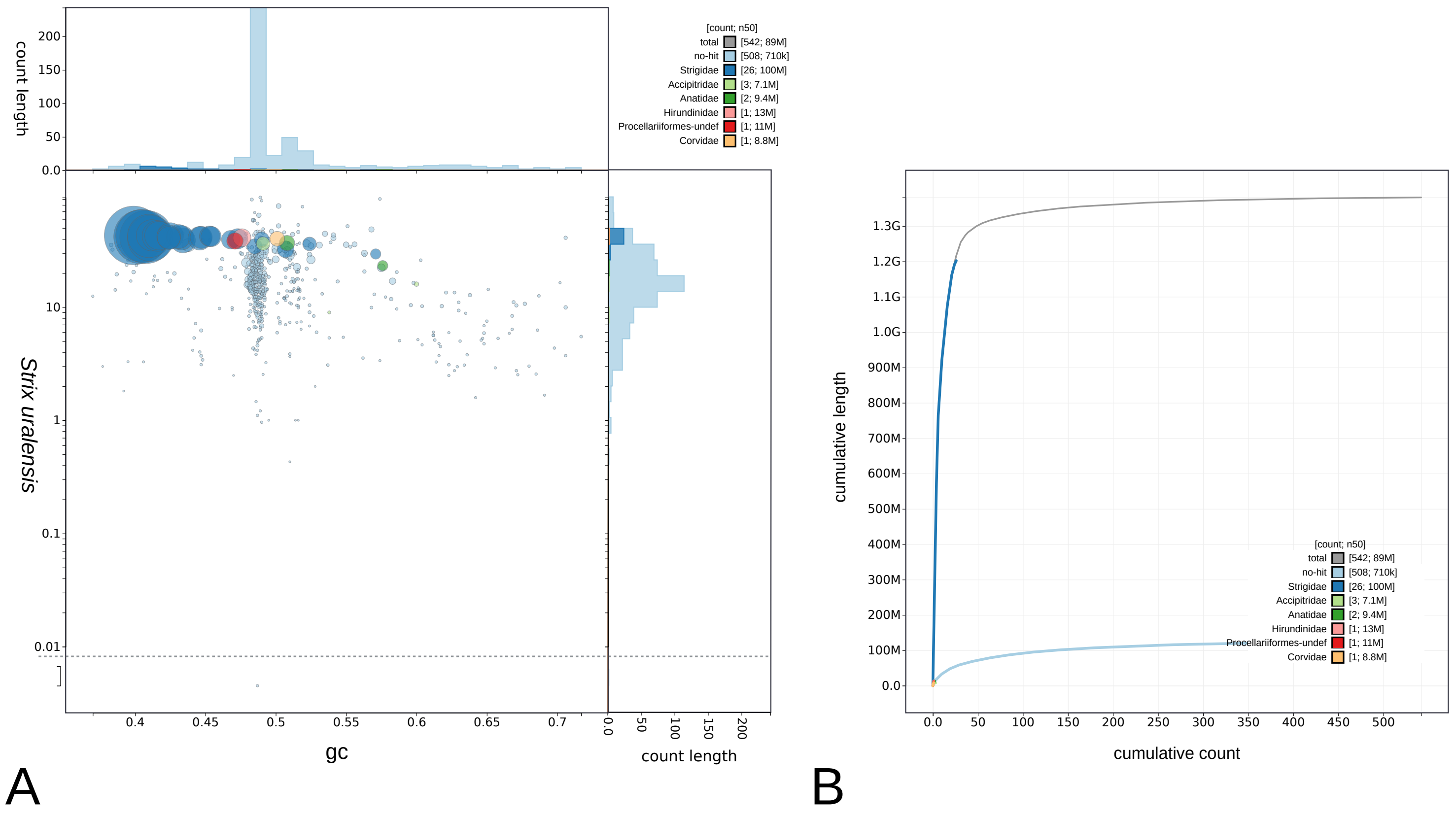


Supplementary Figure S2: Strix uralensis primary haplome BlobToolKit GC-coverage and cumulative sequence plots. A) Blob plot of base coverage in S. uralensis against GC proportion for sequences in S. uralensis primary haplome. Sequences are coloured by phylum. Circles are sized in proportion to sequence length. Histograms show the distribution of sequence length sum along each axis. B) Cumulative sequence length for S. uralensis primary assembly. The grey line shows cumulative length for all sequences. Coloured lines show cumulative lengths of sequences assigned to each phylum using the buscogenes taxrule.

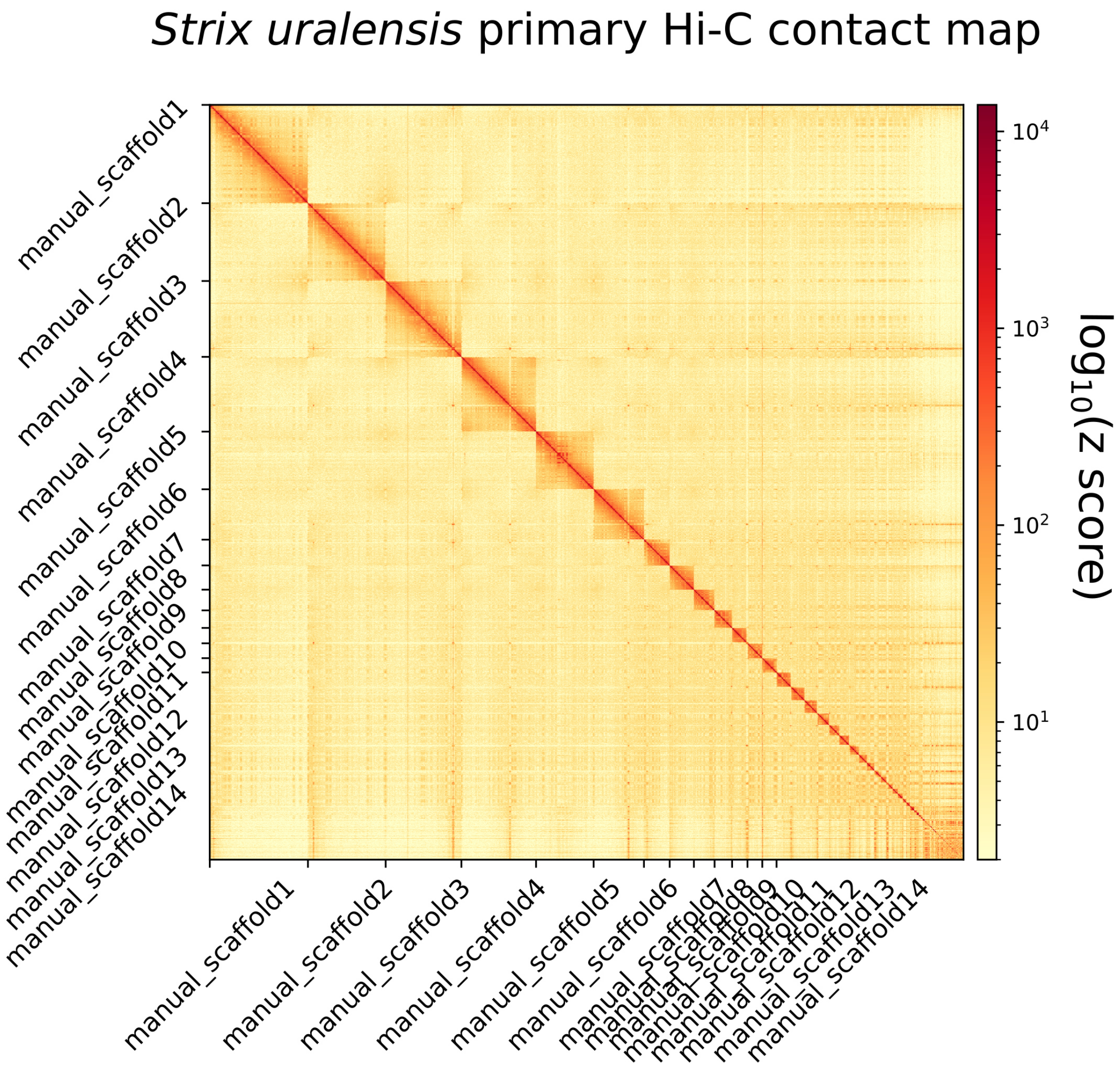


Supplementary Figure S3: *Strix uralensis* primary haplome Hi-C contact map showing spatial interactions for all chromosomes. Chromosomes are ordered by size from left to right and from top to bottom. The red diagonal corresponds to intra-chromosomal contacts and depicts chromosome boundaries. The frequency of contacts is shown on a logarithmic heatmap scale. Plot generated with HiCExplorer.

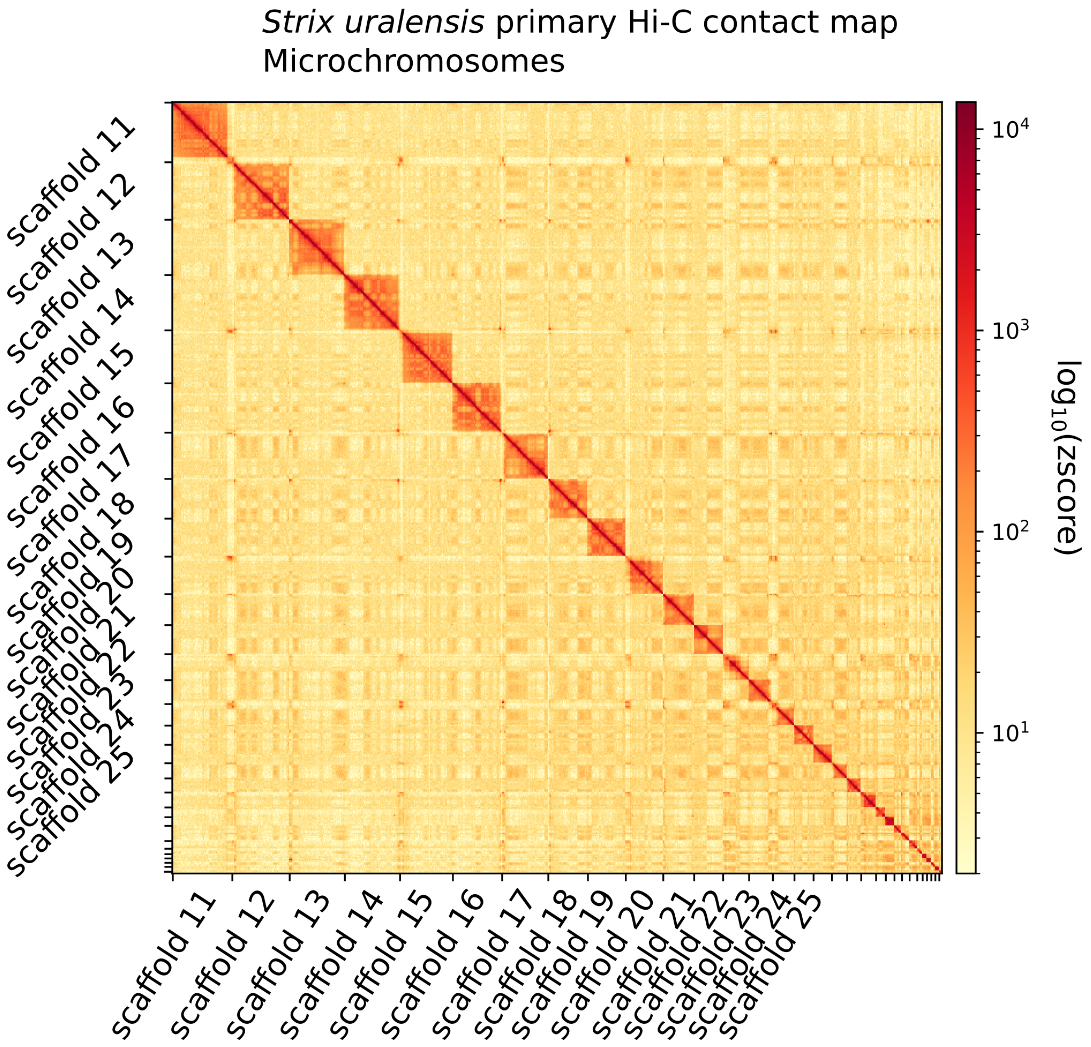


Supplementary Figure S4: *Strix uralensis* primary haplome Hi-C contact map showing spatial interactions between all chromosomes following the ten largest chromosomes. Chromosomes are ordered by size from left to right and from top to bottom. The red diagonal corresponds to intra-chromosomal contacts and depicts chromosome boundaries. The frequency of contacts is shown on a logarithmic heatmap scale. Plot generated with HiCExplorer.

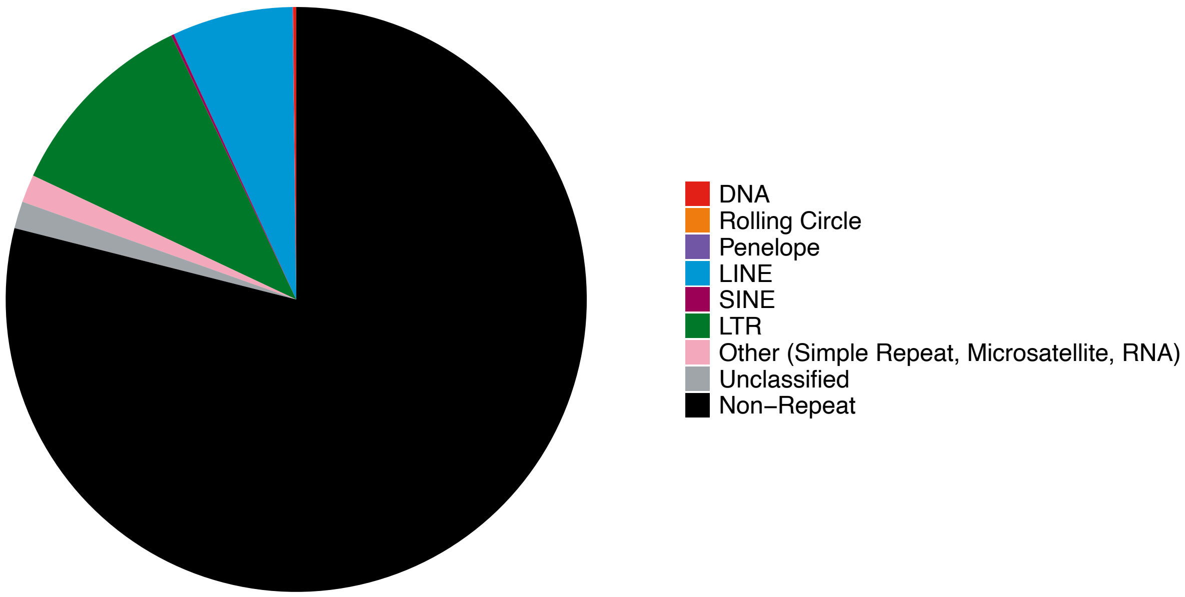


Supplementary Figure S5: Pie chart of repeat family categories estimated by EarlGrey.

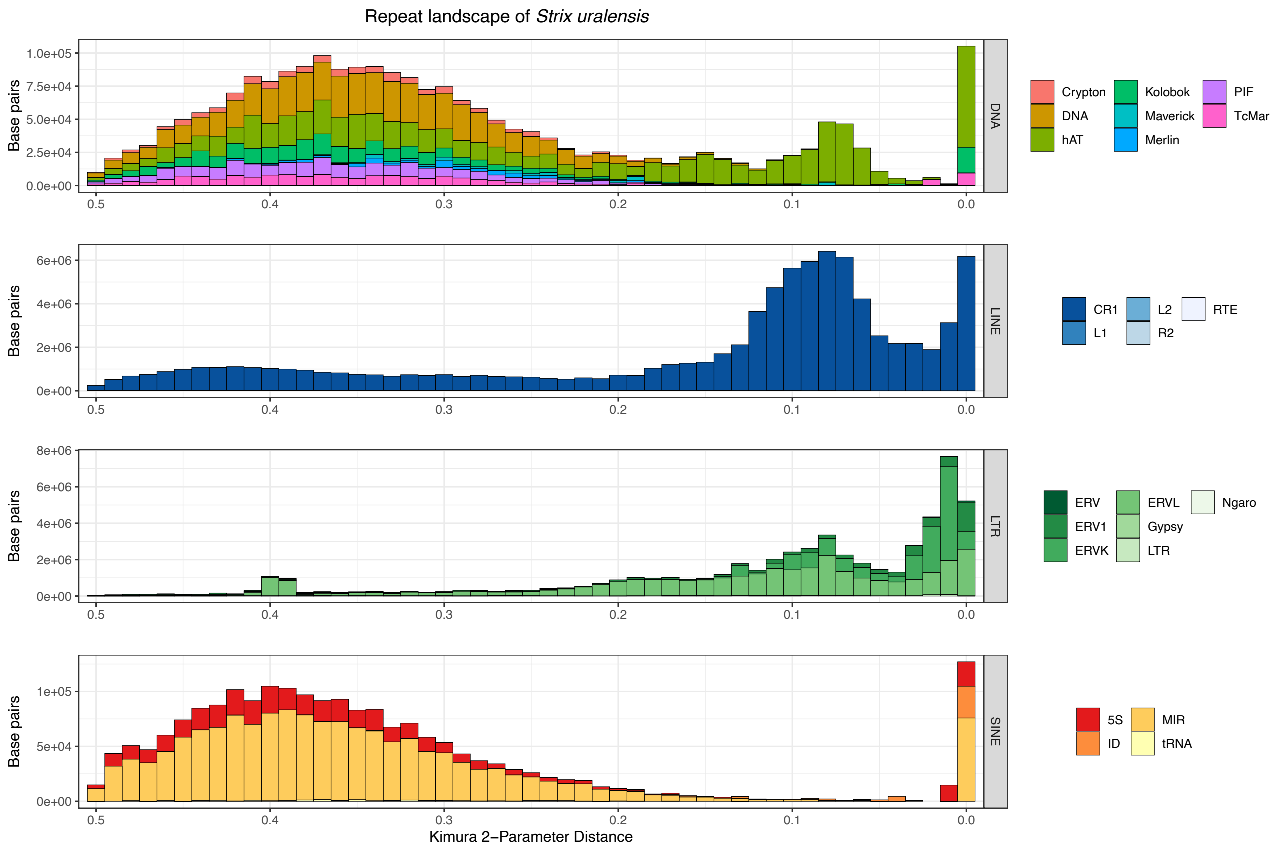


Supplementary Figure S6: Major repeat family categories and subcategories estimated by EarlGrey.

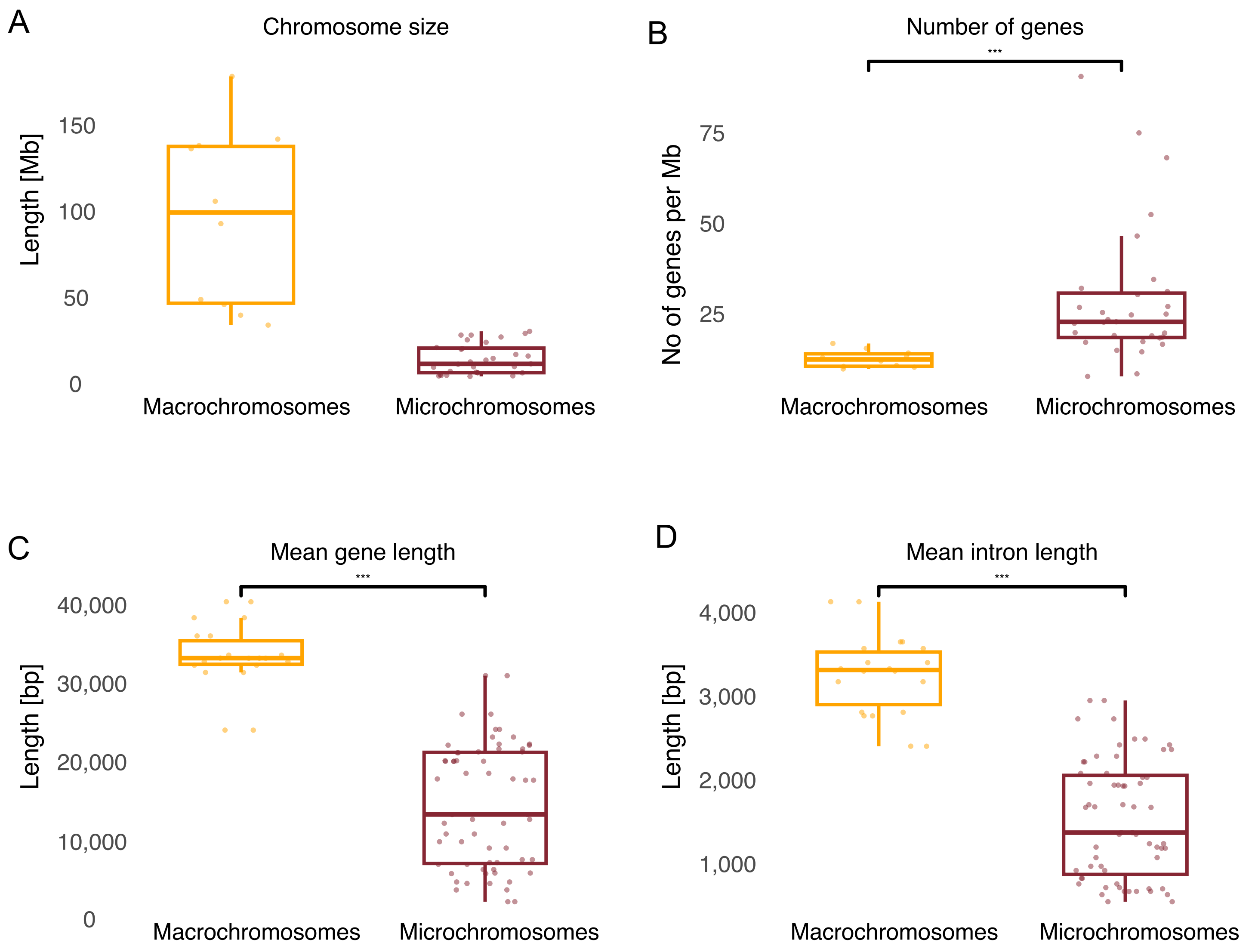


Supplementary Figure S7: Gene structure of the *Strix uralensis* primary haplome. The 41 haploid chromosomes of *Strix uralensis* are divided into macrochromosomes (>30 Mb; n = 10; yellow) and microchromosomes (<30 Mb; n = 31; red). A) Chromosome size distribution of macro- and microchromosomes, B) Gene density of macro- and microchromosomes. C) Mean gene length of macro- and microchromosomes. D) Mean intron length of macro- and microchromosomes. Boxplot centre lines represent the median, box limits the upper and lower quartiles and whiskers the 1.5× interquartile range. Differences were assessed using the Wilcoxon test (*** = p ≤ 0.001).

### GO Terms of gene families unique to *Strix uralensis*

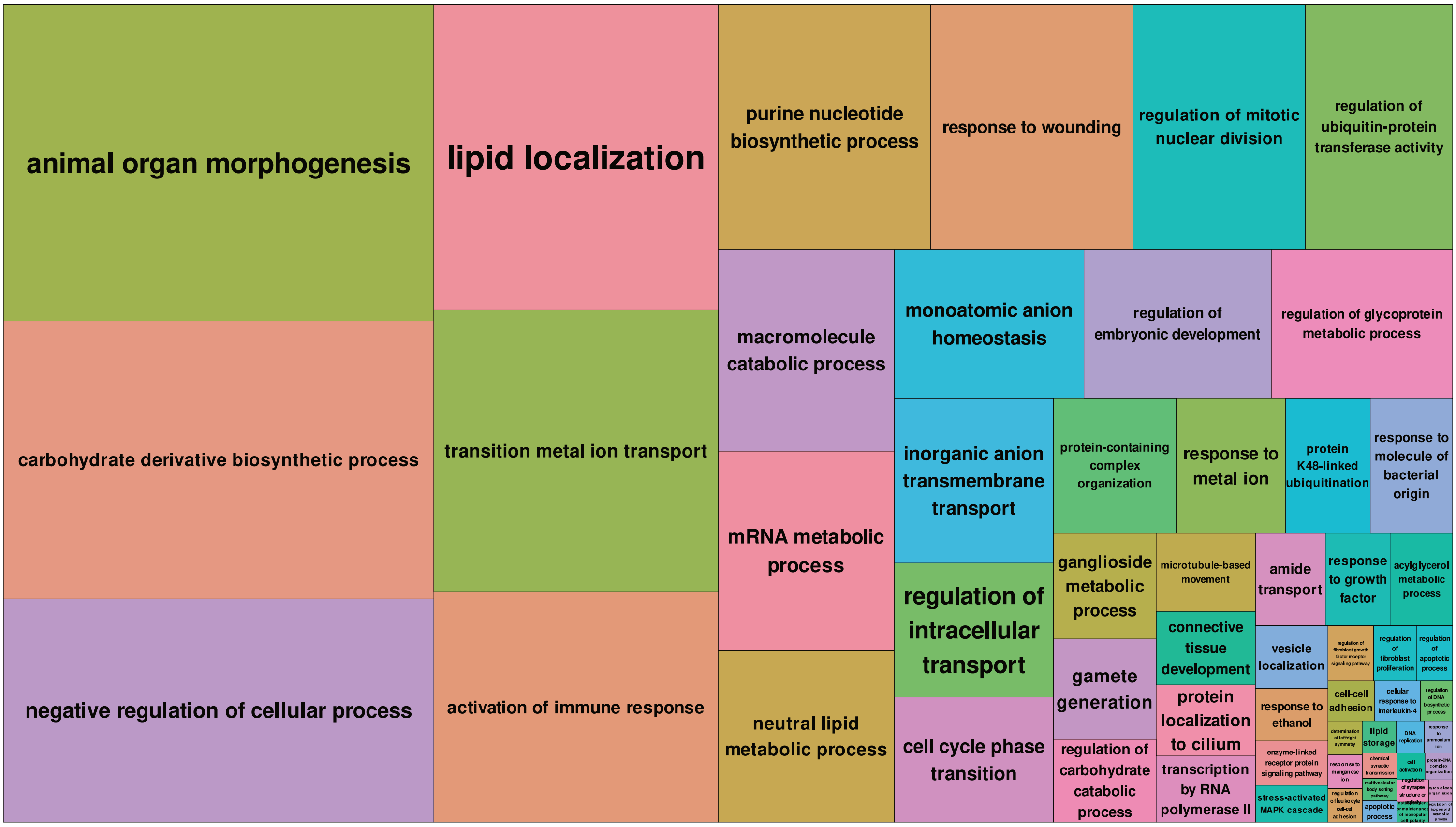


Supplementary Figure S8: Treemap plot of most frequent GO term categories of gene families unique to the *Strix uralensis* primary haplome. Colour of sections is unique to each category and the size scales positively with the GO term frequency.

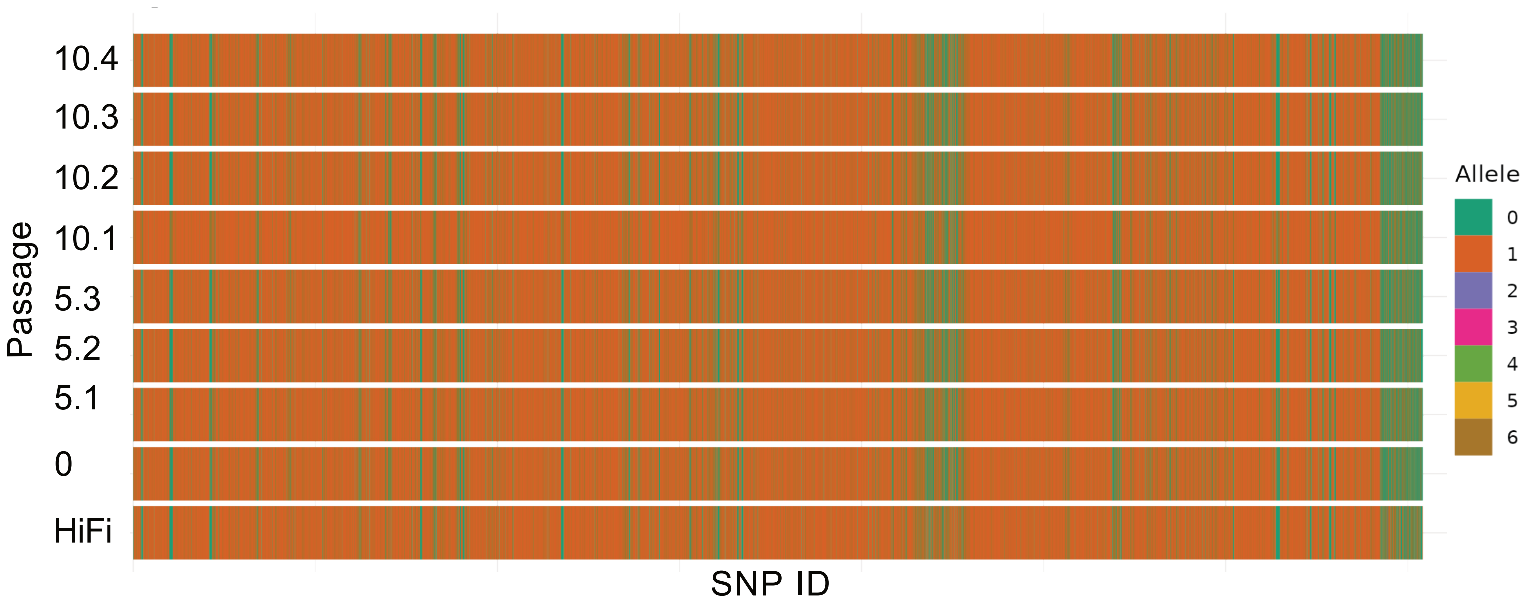


Supplementary Figure S9: Variant calls of each passage. SNPs are ordered by genome position as derived from the variant filed, colour indicates allele as illustrated in the inset and referring to "0|0" = 0, "0|1" = 1, "1|0" = 1, "1|1" = 2, "0|2" = 3, "2|0" = 3, "1|2" = 4, "2|1" = 4, "2|2" = 5, "0|3" = 6, "3|0" = 6.

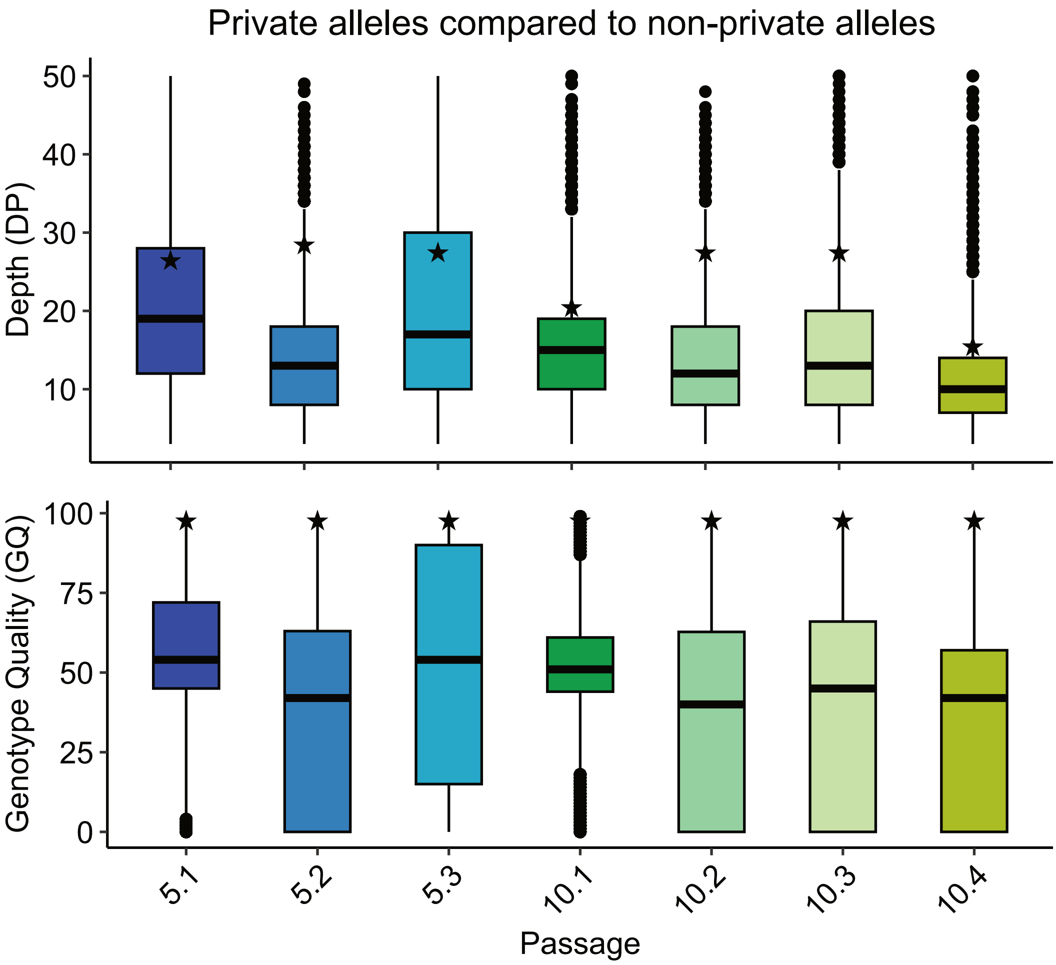


Supplementary Figure S10: Quality assessment of private sites. Median depth (top) and median genotype quality (bottom) of private SNPs in each section compared to non-private SNPs (star).
